# Dawnn: single-cell differential abundance with neural networks for common and rare cell populations

**DOI:** 10.1101/2023.05.05.539427

**Authors:** George T. Hall, Sergi Castellano

## Abstract

A common task in single-cell transcriptomics is the identification of cell populations, often defined by their cell state or type, that are more abundant in one sample or condition than another. Existing methods, however, suffer from either low discovery rates or high rates of false positives, particularly when differences in abundance are confined to some cell states or types or stemming from rare cell populations. We introduce Dawnn, a pre-trained neural network able to find differential abundance with higher accuracy than current tools, both on simulated and biological datasets. Further, Dawnn recovers and extends published findings and discovers more cells in regions with localised differential abundance than existing methods for both common and rare cell types, including extremely uncommon ones such as pulmonary ionocytes. We thus advance the identification of abundance changes in cell populations, regardless of how prevalent the differences and populations are, promising novel biological insights at single-cell resolution.

## 1 Introduction

Single-cell RNA sequencing allows the comparison of samples using the transcriptomes of individual cells. A common task is to identify populations of cells whose states or types exhibit *differential abundance* (DA) across samples or conditions, for example when enriched or depleted in disease, between treated and control conditions in a drug trial, or between patients under the same treatment or treatment time points.

Despite its importance, the identification of cell populations with DA remains a challenge, particularly when it is only found in certain cell states or types or confined to populations that are rare in a sample. This is an important obstacle in biology as individual cell states or types, even if uncommon, often underscore disease or the response to the conditions assessed.

A naïve approach to identifying DA is to create clusters, assume that they correspond to distinct and biologically meaningful populations of cells, and identify those dominated by just one cell state or type across samples or conditions. However, this method can perform poorly if DA is split across multiple clusters, since the signal of enrichment or depletion is split among similar cell populations and therefore becomes challenging to detect.

To address this problem, established methods such as Milo [1], DA-seq [2], and CNA [3] attempt to identify DA without performing clustering whilst moderating the false discovery rate (FDR) [1]. For example, relevant cell populations can be continuously defined and tested across overlapping graph neighborhoods [1]. How well these approaches extend to rare populations is, however, unclear.

In this paper, we present *Dawnn* (*differential abundance with neural networks*), which improves DA identification over existing methods, even for less common cell populations, whilst matching their low FDRs. Dawnn uses a neural network model that has been trained to estimate the relative abundance of each sample or condition in a cell’s neighbourhood. Our approach differs from existing ones since, instead of fitting traditional statistical models to a nearest-neighbour graph of cell labels, we have instead trained Dawnn to estimate the probability with which each cell was drawn from a given sample or condition using simulated training datasets.

Before benchmarking Dawnn, we first create a rigorous formulation of two conflicting definitions of differential abundance in the literature. One definition – that a population of cells, however defined, exhibits DA for a label if that label’s proportion within it differs significantly from its proportion globally – is implicit in some methods but may yield counter-intuitive conclusions in some cases. An alternative definition – which targets DA in local cell populations by testing for an uneven split in labels – is implicit in others. Some methods operate in between these two settings. We discuss the scenarios in which each is appropriate and how they are complementary. We explicitly search for both types of DA with Dawnn.

With these definitions in hand, we evaluate Dawnn’s performance using the bench-marking procedures and datasets introduced by Dann *et al.* to assess Milo [1], comprising simulated and biological datasets of varying complexity. We extend our benchmarking to three additional published real-world datasets. We find that Dawnn yields higher true positive rates than existing methods whilst maintaining low rates of false discovery (the proportion of cells incorrectly claimed to be in populations with DA). Further, Dawnn maintains its good performance on both common and rare cell populations, particularly when searching for DA under the local definition.

We also demonstrate that, as shown by Dann *et al.* for Milo [1], Dawnn is capable of recovering and extending published findings of differential abundance when comparing cirrhotic liver against healthy [4], indicating that its good benchmarking performance likely generalises to real data. Additionally, Dawnn uniquely recovers flow cytometry findings on ionocytes, an extremely rare cell type involved in a number of severe pulmonary pathologies [5], with best performance yielded by its local variant.

Thus, *Dawnn* identifies cell populations with DA between samples or conditions better than existing state-of-the-art algorithms, maintaining a low FDR and promising novel biological insights. Dawnn is available as an R package on CRAN^1^.

## 2 Results

### 2.1 A global view of differential abundance leads to false positives in rare cell populations

Consider an experiment where cells from a patient are treated with a therapy and compared to control (untreated) cells from the same patient (i.e. each cell is labelled either ‘treated’ or ‘control’). Intuitively, a population of distinct cells, however defined, exhibits DA for a treated/control label if it is dominated by that label. For example, a cell state or type that proliferates under treatment may be dominated by treated cells. However, it is unclear how this notion of domination should be defined, leading to conflicting approaches in the literature. Here, we make explicit the differences between the two definitions and when each might be informative.

Under one definition, which we name *global* DA (GDA) and is used in e.g. DA-seq [2], a label is considered to dominate a population of cells if its proportion within that population is significantly higher than its proportion globally. Despite feeling intuitive, the cell populations exhibiting DA under this definition do not necessarily align with the population of cells impacted in an experiment. For example, in Fig. 1, a common cell population that further proliferates following treatment (in red) does not exhibit GDA, whereas the rarer populations (in blue and green) do, despite the treatment not changing their abundance. This limitation is the result of GDA not being determined only by changes in abundance within the population of interest, and leads to false positives in rare cell populations.

**Fig. 1:**
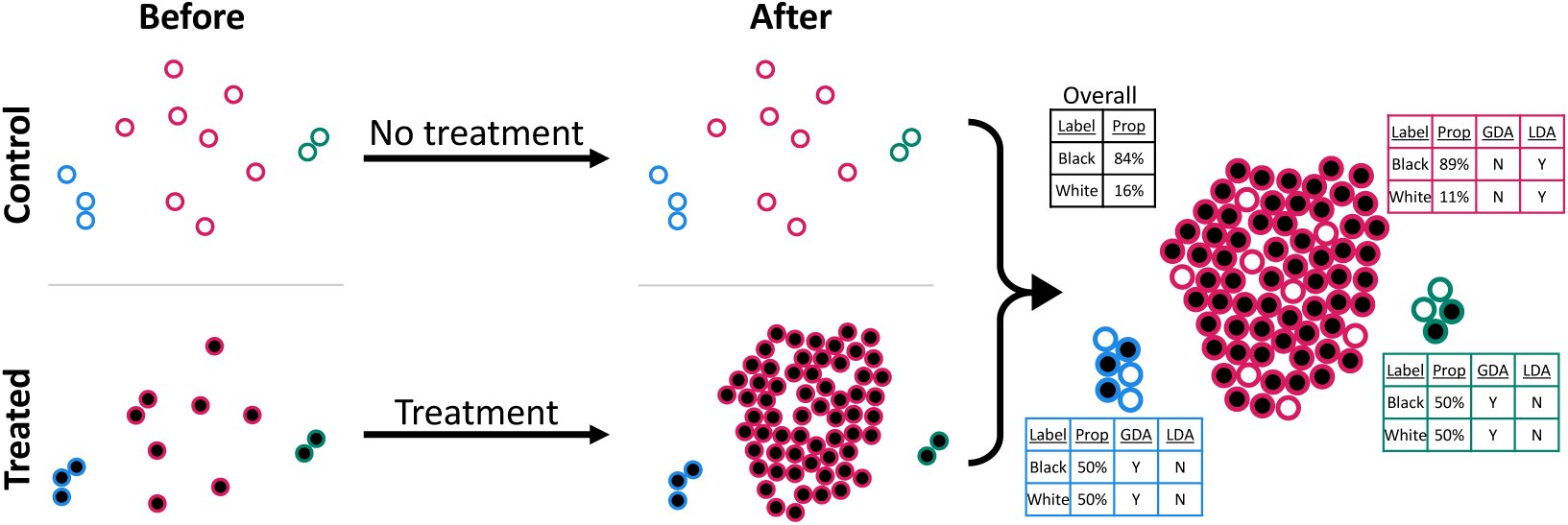
A scenario where the cell populations exhibiting GDA, before and after treatment, do not align with those changing in abundance. The three cell types have different outline colours. Control cells are filled with white and treated cells with black. Treated cells from the red cell type proliferate with treatment and exhibit DA under LDA but not GDA.

A *local* definition of DA (*LDA*) can address these shortcomings. In contrast to GDA, under LDA a label is considered to dominate a population of cells if its proportion within that population differs significantly from 0.5. In the above example, LDA identifies that only the red cell type increases in abundance following treatment (Fig. 1). We believe this definition to be more intuitive and helpful unless the two samples or conditions compared have large differences in their number of cells, which could result from uncontrolled technical biases (e.g. decreased capture efficiency in droplets for cells in one sample or condition or differences across samples in their preservation or processing methods). We do not address these experimental biases in this work or discuss how to control for them, though we have offered some technical guidance for challenging tissues in [6].

We provide more rigorous definitions of local and global DA and investigate how each behaves in different scenarios in Section 5 in Supplementary Information.

### 2.2 Dawnn detects differential abundance using a pre-trained neural network

Dawnn is designed to identify differential abundance in cell populations comprising two samples or conditions. Whilst it differs from existing algorithms in its methodology, it employs the same basic assumptions, namely that each cell is assigned one of two labels (*S_a_* or *S_b_*) and is associated with an individual unknown probability *p* of having the label *S_a_* (in effect, of being drawn from the sample or condition associated with label *S_a_*). We denote as 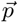 the vector of all probabilities *p* associated with a dataset. A given single-cell transcriptomic dataset is thus considered an instantiation of labellings generated according to 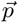. Under this model, the task of detecting populations of cells with DA is equivalent to that of identifying cells for which *p* differs substantially from a null distribution that assumes no DA, for a given definition of DA. Dawnn searches for both LDA and GDA to provide the user with as much information as possible and to aid diagnosis of experimental issues (see Discussion).

In general, DA detection methods estimate 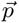 from a distribution of sample labels on a *K*-nearest neighbour graph. Batch effects can be mitigated by generating the KNN from a batch-corrected dimensionality reduction (e.g. Harmony [7]). Existing DA detection algorithms estimate 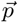 by fitting a model (e.g. Milo’s generalised linear or DA-seq’s logistic regression models) to this distribution. Dawnn, on the other hand, employs a neural network (Section 11.1 in Supplementary Information) that has been trained to estimate 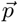 using simulated training data. Dawnn first estimates 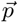 from the sample labels of the 1,000 nearest neighbours of each cell, then generates a null distribution of labels to represent a dataset with no differential abundance (in terms of either local or global differential abundance). It then estimates 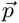 for this null distribution dataset. For each cell, Dawnn tests the null hypothesis that it is in a population of cells with no DA, calculating a p-value by fitting a beta distribution to the estimates of 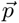 from the null distribution data and calculating the probability of observing at least such an extreme a *p* under this distribution. Dawnn controls the false discovery rate (FDR) – the proportion of cells incorrectly classified as belonging to regions exhibiting DA – using the Benjamini–Yekutieli procedure [8], a variant of the Benjamini-Hochberg procedure [9] that does not assume independence between hypotheses. This more conservative FDR-controlling mechanism is necessary since the hypotheses associated with neighbouring cells will be correlated. Pseudocode for Dawnn is given in Algorithm 1.

#### Algorithm 1: Dawnn (*Diflerential abundance with neural networks*)

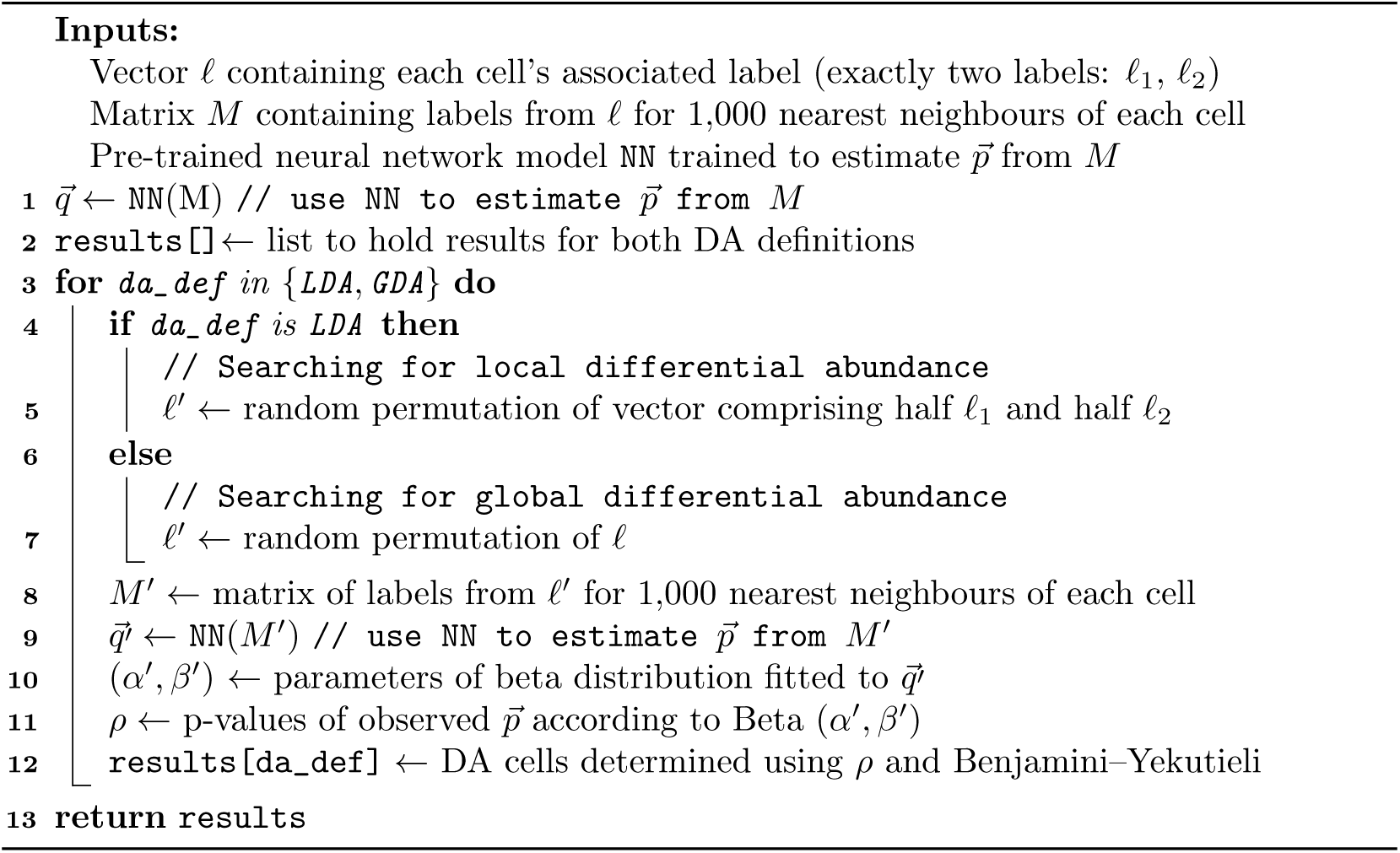

We selected the architecture of Dawnn’s neural network from a pool of candidates assessed using *K*-fold cross validation (Section 11.2 in Supplementary Information). Our algorithm training and selection procedures were designed to yield an unbiased model capable of estimating 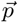 across many real-world datasets. To maximise generalisation and avoid bias during benchmarking, we designed Dawnn’s training set to make no assumptions about the overall structure of cell populations and only weak assumptions about the relationship between the labels of cells within each neighbourhood (Section 4.1.1).

### 2.3 Dawnn detects regions exhibiting DA with higher accuracy than existing methods

We benchmarked Dawnn against Milo and DA-seq, since these were the two best-performing approaches in Dann *et al.* [1]. We also benchmarked against CNA, which was developed concurrently with Milo and therefore omitted from [1]. We follow the benchmarking of Dann *et al.* (Section 4.2.2), creating simulated and real datasets using their described approach [1]. We further assessed real-world performance using three additional biological datasets: one from a publication of ours examining the effects of accelerating skin sheet growth [10]; another from a publication characterising organoids of bile ducts [11]; and a large dataset comprising cells from the heart in a study of congenital heart disease [12]. Table 1 summarises our benchmarking datasets.

**Table 1:** Test datasets used to assess true positive and false discovery rates.

| Dataset | Source | Number of cells | Number of cell types |
| --- | --- | --- | --- |
| Discrete clusters | Simulation [1] | 2,700 | NA |
| Linear trajectory | Simulation [1] | 10,000 | NA |
| Branching trajectory | Simulation [1] | 7,500 | NA |
| Mouse gastrulation | Experiment [13] | 64,018 | 37 |
| Accelerated keratinocytes | Experiment [10] | 18,787 | 3 |
| Bile duct organoids | Experiment [11] | 72,803 | 1 |
| Congenital heart disease | Experiment [12] | 157,293 | 14 |

The simulated and experimental datasets provide the structure of cell populations to which simulated labels can be assigned, providing the ground truth necessary to measure algorithm performance.

Six cell populations were created for each simulated dataset. For each population, we simulated multiple distributions of sample labels, with each label distribution defined by the subpopulation exhibiting differential abundance and the extent of differential abundance in this subpopulation. For each choice of differentially abundant subpopulation, three sets of sample labels were simulated for each maximum log_2_-fold change in the set {1.22, 1.58, 2, 2.5, 3.17, 4.25} (the same log_2_-fold changes employed by Dann *et al.* [1]) We then formed ground truth labels for LDA and GDA.

We used a similar methodology to simulate sample labels for datasets from real experiments. Here, in each dataset, eight subpopulations were chosen to be differentially abundant. These subpopulations corresponded to annotated cell types in the mouse gastrulation dataset and to clusters in the keratinocyte, organoid, and heart datasets. The maximum log-fold changes were identical to those in the simulated datasets.

We assessed algorithm performance by calculating the true positive rate (TPR) – the proportion of cells correctly identified as being in a DA region – and false discovery rate (FDR) – the proportion of cells incorrectly classified as belonging to a DA region. Dawnn and Milo allow the specification of a maximum acceptable FDR. We set these thresholds to 10%, as in Dann *et al.* [1].

We carried out separate benchmarking for the detection of LDA and GDA. Since we consider LDA to be the more intuitive definition of DA (Section 2.1 and Discussion), we primarily assess the performance of algorithms under this definition. For completeness, since it may sometimes be useful to detect populations exhibiting GDA, we include benchmarking for GDA detection (Section 6 in Supplementary Information). Although Dawnn searches for both LDA and GDA concurrently, for clarity, we treat these complementary outputs as two separate algorithms in our results (‘Dawnn-L’ and ‘Dawnn-G’), respectively.

#### Simulated datasets

##### Discrete clusters

For local DA (Fig. 2a), both Dawnn methods and CNA had high TPRs. However, only Dawnn-L maintained FDRs consistently below the 10% target. For global DA (Supplementary Fig. 2a), all methods were able to detect cells in DA regions. Dawnn-G and CNA had the highest TPRs. All FDRs were close to 0.

**Fig. 2:**
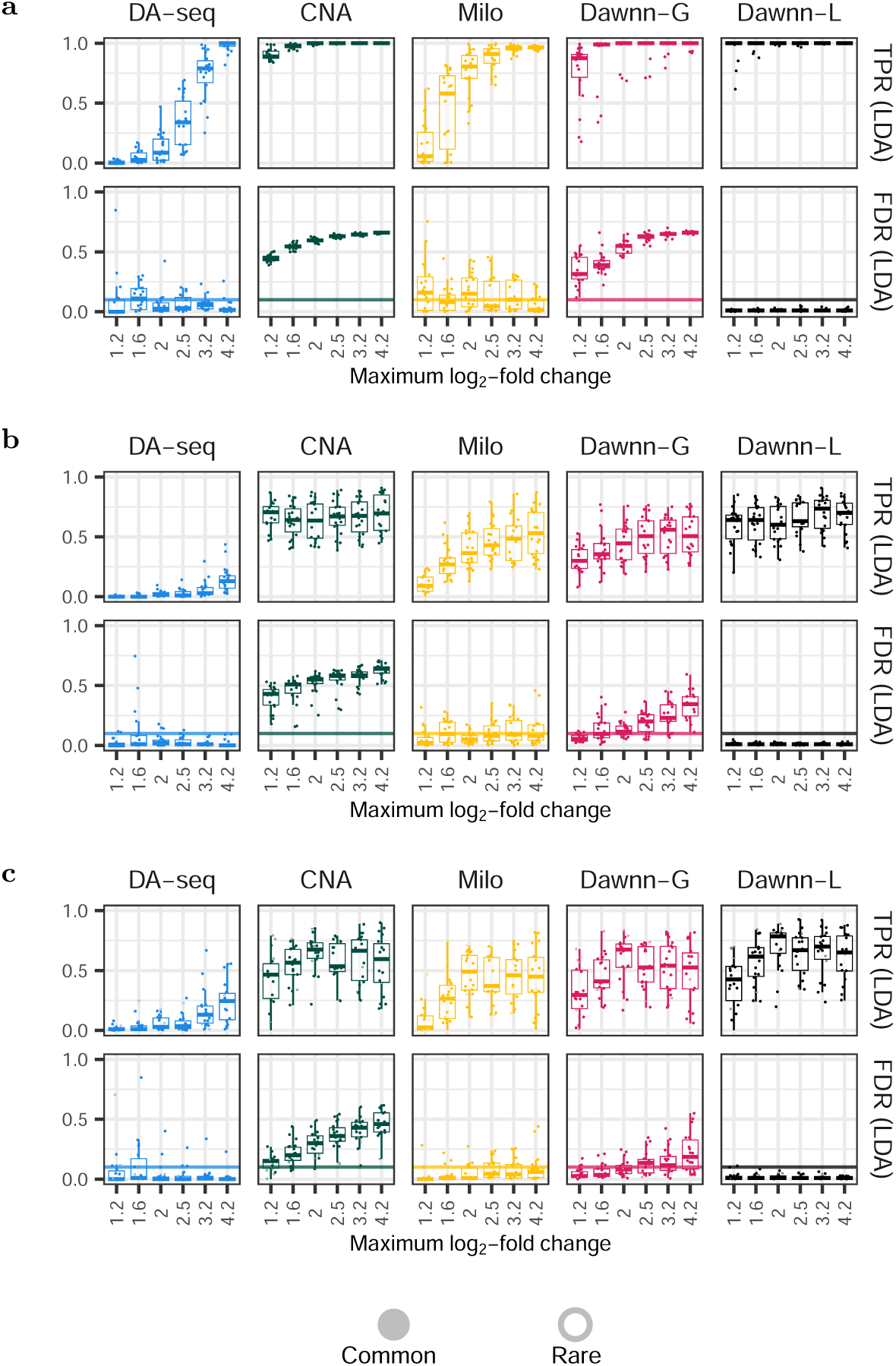
True positive and false discovery rates on simulated datasets for LDA. **a** discrete clusters; **b** linear trajectories; **c** branching trajectories. Filled circles correspond to common cell types (>5% abundance); white circles to rare (*≤*5%). Dawnn-L is the only algorithm able to identify LDA cells without high FDR.

##### Linear trajectories

For local DA (Fig. 2b), all methods except DA-seq had moderate TPRs. All methods except Dawnn-L had FDRs above 10%, with CNA and Dawnn-G exhibiting the highest. For global DA (Supplementary Fig. 2b), CNA had highest TPRs but also high FDRs, only falling below 10% for high LFCs. Dawnn-G and Milo had similar performances, with medium TPRs and FDRs below 10%. Whilst Dawnn-L had similarly high TPRs, it also had FDRs above 10%.

##### Branching trajectories

For local DA (Fig. 2c), all methods except DA-seq achieved moderate TPRs but all except Dawnn-L also failed to moderate FDRs below 10%. For global DA (Supplementary Fig. 2c), CNA and both Dawnn variants achieved high TPRs, although all methods (especially CNA) failed to consistently moderate FDRs below 10%.

#### Experimental datasets

##### Mouse gastrulation

For LDA (Fig. 3a), all methods again had moderate TPRs, although DA-seq only for higher LFCs. The FDRs of Dawnn-G and CNA often exceeded 10%. For GDA (Supplementary Fig. 3a), all methods had moderate TPRs (with CNA the highest), although DA-seq again only for higher LFCs. CNA and, to a lesser extent Dawnn-L and Dawnn-G, had FDRs exceeding the 10% threshold.

**Fig. 3:**
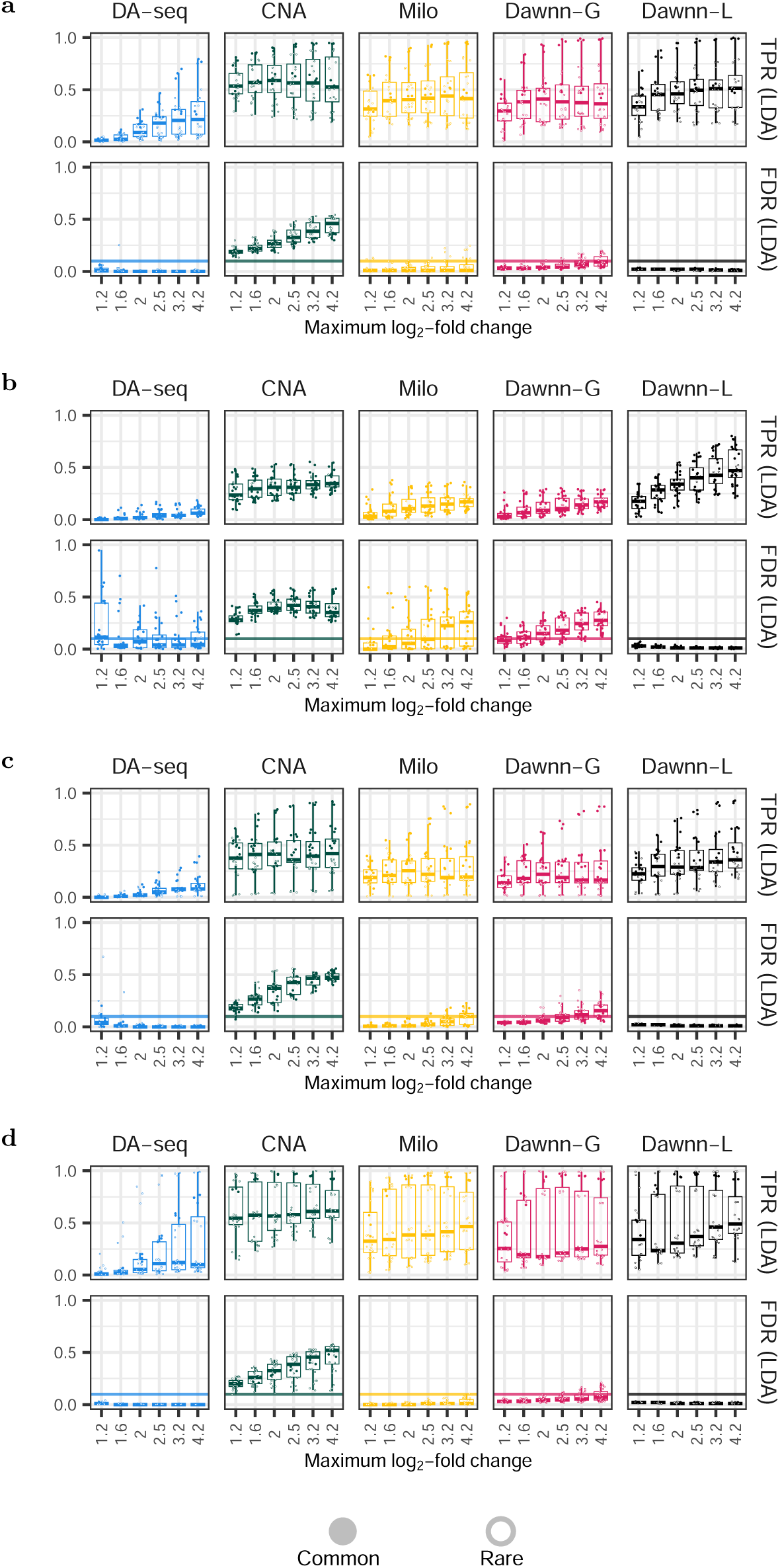
True positive and false discovery rates on simulated datasets for LDA. **a** mouse embryo; **b** keratinocytes; **c** organoids; **d** heart. Filled circles correspond to common cell types (>5% abundance); white circles to rare (*≤*5%). Dawnn-L is the only method that can reliably control its FDRs below 10%.

##### Growth-accelerated keratinocytes

For LDA (Fig. 3b), only CNA and Dawnn-L exhibited high TPRs. All methods except Dawnn-L failed to moderate FDRs below 10%. For GDA (Supplementary Fig. 3b), all methods except DA-seq had moderate TPRs (especially for high FDRs), with CNA exhibiting the highest. However, all methods had FDRs over the 10% threshold. For larger LFCs, the mean FDRs of Milo, DA-seq and Dawnn-G dropped below 10%.

##### Bile duct organoids

For LDA (Fig. 3c), all methods except DA-seq had moderate TPRs. Only Dawnn-L kept FDRs below 10%. For GDA (Supplementary Fig. 3c), all methods had low TPRs, with CNA and Dawnn-L the highest. All methods generally had FDRs exceeding 10%, although those of all methods except DA-seq dropped below 10% for larger LFCs.

##### Heart cells

For LDA (Fig. 3d), CNA had the highest TPRs, followed by Milo and Dawnn-L (which were comparable). The TPRs of Dawnn-G were slightly lower and those of DA-seq were much lower. DA-seq, Milo and Dawnn-L all moderated FDRs below 10%, whilst those of Dawnn-G slightly exceeded this target and CNA did so greatly. For GDA (Supplementary Fig. 3d), all methods except DA-seq had moderate TPRs, with CNA the highest, followed by Milo and Dawnn-L. All methods except CNA had mean FDRs below 10%.

#### 2.3.1 Results summary

For LDA, all methods except DA-seq exhibited similar TPRs, with Dawnn-L and CNA often having the highest. However, Dawnn-L was the only method where FDRs were consistently under 10%.

For GDA, all methods generally had high TPRs. CNA usually had the highest, followed by Dawnn-G. TPRs decreased as datasets became more complex. However, the FDRs of CNA often exceeded 10%, rendering its outputs unreliable. Dawnn-G was the method with highest TPR that kept FDRs generally under 10%, although at times this threshold was exceeded.

We validated these conclusions with statistical analyses (Section 7 in Supplementary Information), supporting our above conclusions.

### 2.4 Dawnn maintains accuracy for rare cell populations

To better understand the influence of cell type rarity on the performance of DA detection methods, we displayed the benchmarking results above by points coloured by whether the DA is driven by a common (>5% abundance) cell type or a rare (≤ 5%) one. For datasets with both common and rare populations (i.e. branching trajectories and all experimental datasets), we entirely separate the plots for these two categories of DA in Supplementary Fig. 5. In Supplementary Figs. 6 and 7, we also plot TPRs and FDRs in relation to the rarity of the differentially abundance cell population for all datasets.

In general, DA contained within rarer cell populations was more difficult for all methods to detect (i.e. TPRs were lower). Further, these cases also resulted in higher FDRs, meaning that other, non-DA cell populations were more likely to be assessed as DA if the DA is contained within a rare cell type, as expected (Fig. 1). When searching for global differential abundance, the FDRs of all methods frequently exceeded the 10% specified threshold when DA was contained within a rare cell population. When searching for local differential abundance, however, Dawnn-L was able to maintain FDRs below 10%, both for common and rare cell populations. All other methods, conversely, often exceeded this threshold for DA driven by rare cell types. When searching for local DA, therefore, the estimates of Dawnn-L are more reliable than those of the other methods.

### 2.5 Dawnn makes biologically plausible discoveries

We ran Dawnn, DA-seq and Milo on biological datasets to investigate which methods are able to recover published findings. We excluded CNA since its high rates of false discovery (Section 2.3) mean that labelling more cells as in regions of DA is not necessarily desirable.

Each method exhibited a clear threshold in the log-fold change (LFC) of sample abundance in a neighbourhood (Milo) or cell’s neighbourhood (all other methods) before calling a neighbourhood / cell as DA. This threshold varied between local and global methods. Since, in global methods (DA-seq and Dawnn-G), the null distribution assumes the same sample abundances as in the original data, this threshold was centred at the LFC corresponding to the global ratio between the two samples. Consequently, in datasets where one sample is much more abundant than the other, cells could be called as being in a region dominated by the less abundant sample even if the other sample was still more abundant. For example, a cell in a region with an LFC of −1 could be labelled as in a region dominated by the sample associated with positive LFC if the global LFC is −4. In Dawnn-L, on the other hand, the DA-calling LFC threshold was centred at 0, since the null distribution assumes equal proportions of both samples. In both datasets, the size of the interval of LFCs for which neighbourhoods / cells were not called as DA was smaller in Dawnn compared to Milo and DA-seq, yielding Dawnn’s higher discovery rates.

#### 2.5.1 Liver cirrhosis

Dann *et al.* [1] demonstrate that Milo is capable of recovering findings from Ramachandran *et al.*, who compared samples of healthy and cirrhotic livers [4]. Both Dawnn and DA-seq were able to recover these findings at the single-cell level rather than the neighbourhoods operated on by Milo (Fig. 4). Further, Dawnn was capable of identifying regions of differential abundance that are missed by DA-seq and Milo due to their lower rates of discovery.

**Fig. 4:**
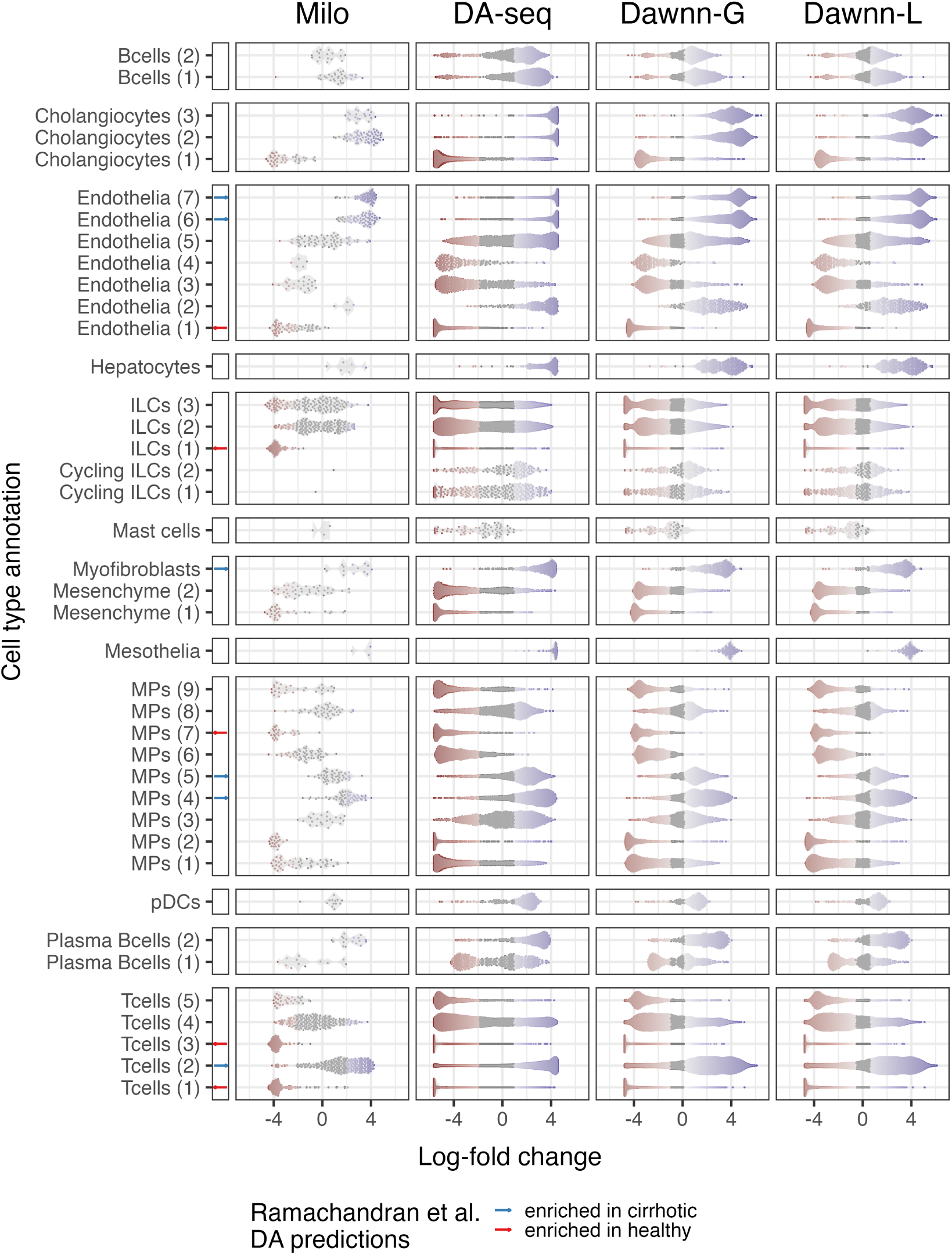
Log-fold changes estimated by Milo, DA-seq, Dawnn-G, and Dawnn-L on the liver cirrhosis dataset of Ramachandran *et al.* [4]. Positive log-fold changes correspond to increased abundance in diseased samples; negative to increased abundance in healthy.

For cell types labelled as differentially abundant by Ramachandran *et al.*, the outputs of all methods report largely similar extents of DA. Within these groups of cells, as Dann *et al.* reported for Milo, both Dawnn and DA-seq identify sub-populations differentially abundant in the opposite direction to the larger population, potentially indicating subpopulations within annotated cell types that are different affected by cirrhosis. For some cell types, such as T Cells (5) and PDCs, the cell-level nature of Dawnn and DA-seq allows them to identify DA in populations where Milo fails to do so.

For almost all cell types, Dawnn labels more cells as belonging to regions exhibiting DA than other approaches, since the log-fold change cutoff for calling DA is approximately ±0.5 rather than the approximately ±1.5 and ±2 in DA-seq and Milo, respectively. This equates to over 2,300 more cells being identified as in regions of DA than are labelled as such by DA-seq (an increase of 5%). Supplementary Table 5 contains the number of cells of each type predicted to be in such region of DA by Dawnn and DA-seq, visualised in Supplementary Fig. 8. Both versions of Dawnn have a higher discovery rate than DA-seq for each cell type, indicating that its ability to identify biological insights missed by existing methods applies to both rare and common populations. Dawnn-L in particular identifies more DA in uncommon cell types compared to global methods. For instance, for the rare Mast cell type, Dawnn-L estimates 20 more cells to be in a region of DA than is done so by DA-seq, an increase of 69%. Even for abundant populations, such as ILCs, Dawnn-L identifies an additional 599 cells as belonging to such a region, an increase of 7% over DA-seq. For the majority of cell types, both Dawnn-L and Dawnn-G estimate at least 4% more cells to be in a region of differential abundance than is done so by DA-seq.

#### 2.5.2 Ionocytes and other rare pulmonary cell types in COPD

We further investigated the ability of Dawnn and other methods to identify DA in uncommon cell populations in another dataset. Ionocytes are a rare cell type in the human airway, accounting for approximately 1% of cells in its epithelium in human and mouse [14, 15]. They are nevertheless important in a number of respiratory diseases, including cystic fibrosis and chronic obstructive pulmonary disease (COPD) due to its high expression of CFTR [14–16]. Importantly, using Barttin (encoded by the BSND gene) as a marker for ionocytes, flow cytometry suggests a two-fold reduction in ionocytes cells in COPD [17]. To assess whether DA methods can reproduce this finding on such extremely rare cell type, as well as other rare pulmonary cell types, we interrogated single-cell transcriptomics profiles of healthy and COPD lungs [5] (Fig. 5).

**Fig. 5:**
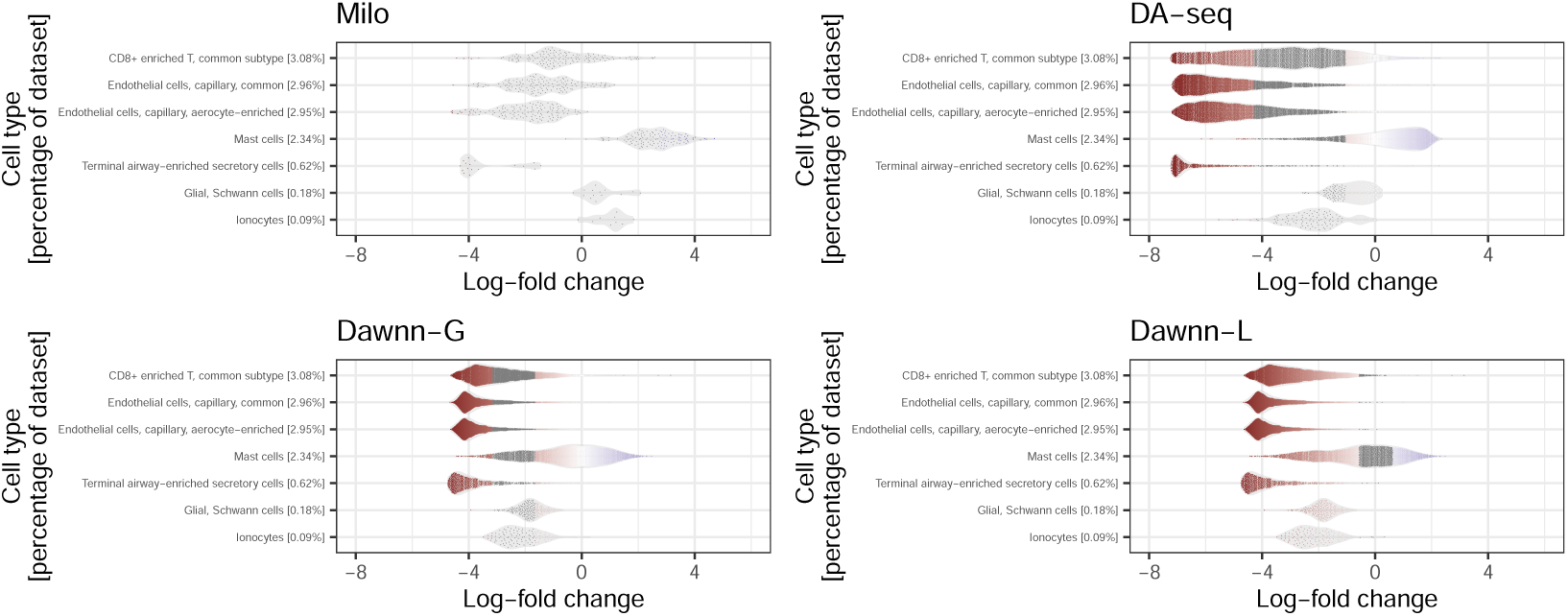
Log-fold changes estimated on a subset of cell types in COPD of Rustam *et al.* [5]. Positive log-fold changes correspond to increased abundance in disease; negative to increased abundance in healthy. Other cell types displayed in Supplementary Fig. 9.

The global methods DA-seq and Dawnn-G both identified small populations of ionocytes more and less abundant in diseased compared to healthy samples (albeit in the order of tens of cells, due to the rarity of the cell type), whilst claiming that the majority exhibited no DA between conditions. Milo’s neighbourhood-based approach created few ionocyte-associated neighbourhoods and did not identify DA in any of them. Dawnn-L, on the other hand, identified that nearly all ionocytes were less abundant in COPD samples, in agreement with published flow cytometry results on this disease [17].

Further, Rustam *et al.* [5] suggested other relatively uncommon cell types with changing abundance in COPD. In particular, Terminal airway-enriched secretory (TASC) and capillary cells (En-c) may be reduced in disease, whereas Glial/Schwann cells (Gli), mast cells (MC) and some lymphoid cells (CD8-T1) may increase. Notably, Milo identified little DA in these cell types (Fig. 5), whereas DA-seq and Dawnn identified more. The global methods, DA-seq and Dawnn-G, identified similar levels of DA in the more abundant cell types (TASC, En-C, MC, CD8-T1). As in Section 2.5.1, Dawnn-G identified a larger proportion of cell types as DA compared to DA-seq. However, in the rare cell types, such as Glial/Schwann cells, global methods only claimed a small number of cells to be in regions of DA. Dawnn-L, however, identified DA in both these cell types, matching the findings of Rustam *et al.* [5].

In Supplementary Fig. 9, we extend the results of Fig. 5 to all cell types. Supplementary Table 6 contains the number of cells of each type predicted to be in such region of DA by Dawnn and DA-seq, visualised in Supplementary Fig. 10.

### 2.6 Dawnn runtime comparable to Milo

We compared the runtime of Dawnn against those of DA-seq, CNA, and Milo on different sizes of datasets (Fig. 6). Importantly, for datasets up to 250,000 cells, all methods except DA-seq ran in below an hour on a standard laptop, making them practical for most studies. The fastest method by far was CNA, followed by Dawnn and Milo, whose runtimes were comparable. DA-seq was the slowest method, so much so that it was not run for the largest dataset. Crucially, the runtime of all methods except DA-seq appears to scale linearly with the number of cells in the dataset, allowing them to be used on large datasets without becoming prohibitively slow.

**Fig. 6:**
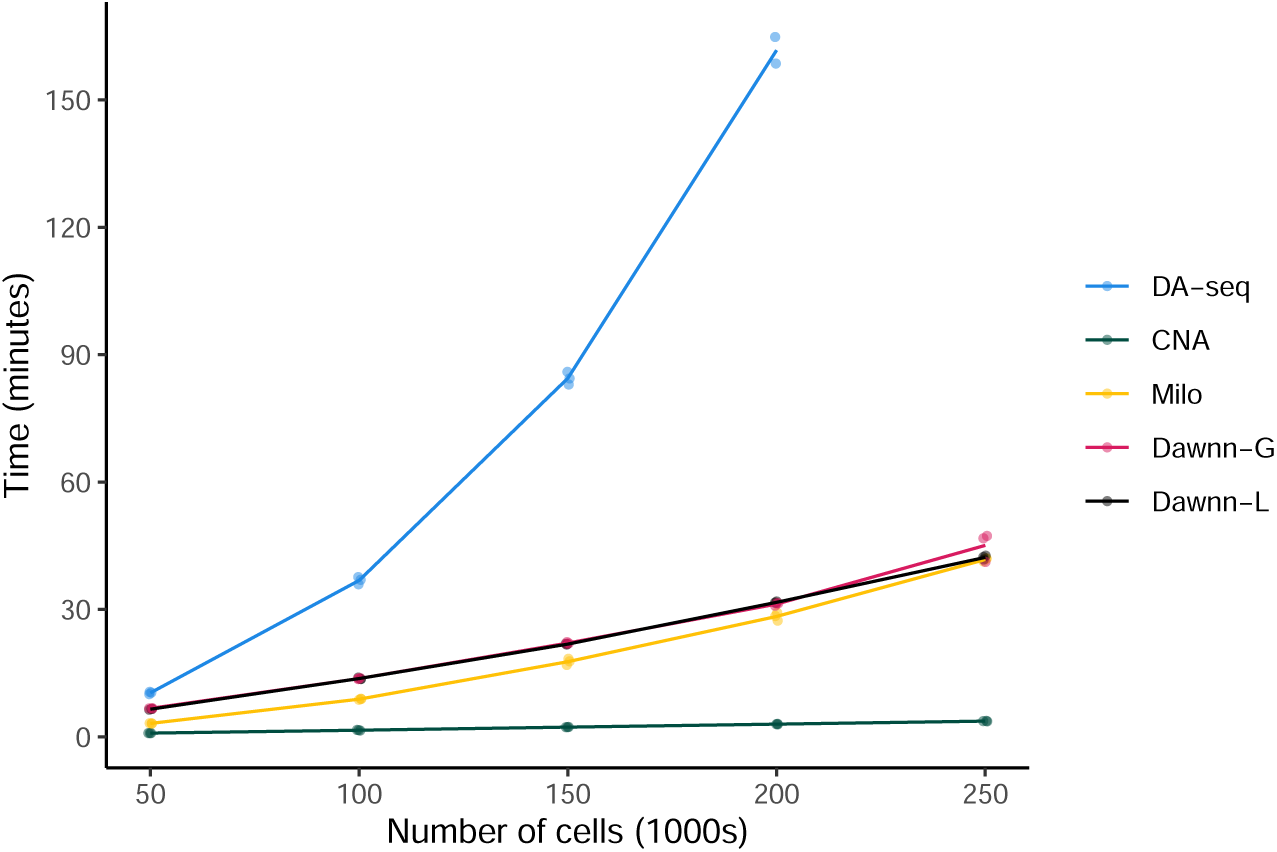
Comparison of runtimes of Dawnn, CNA, DA-seq, and Milo on datasets of increasing size. Dataset size (in terms of thousands of cells) is shown on the x axis, runtime (in minutes) is shown on the y axis.

## 3 Discussion

In single-cell transcriptomics, the identification of cell populations more abundant in one sample or condition than another is a crucial task. However, current methods are unable to do so with high rates of discovery whilst avoiding false positives. DA driven by changing abundance in rare cell types is also often critical but challenging to identify. We first show that existing methods search for different types of DA, one of which may sometimes lead to counter-intuitive results due to its global nature. We claim that an alternative, local, definition may, in some contexts, better align with the cells of interest (i.e. those which are driving the change in abundance). The definition used by some methods – which we term global differential abundance (GDA) – considers a label to exhibit differential abundance in a cell population if its abundance within the population differs significantly from its abundance overall. We show that GDA may yield counter-intuitive results if, for example, the difference in the number of cells with each label is only present in a single, common, cell type, since an unperturbed cell type would still exhibit GDA, and the perturbed one may not. An alternative definition, *local* differential abundance (LDA), considers a label to exhibit differential abundance in a cell population if it is assigned to significantly more than half of the population’s cells. In the same scenario, LDA would only consider the perturbed cell type as exhibiting DA. However, in cases of large differences between the abundance of labels (large class imbalance), LDA will likely yield counter-intuitive results. It is therefore important to minimise sampling and processing biases that change cell proportions [6] and to allow users to explore both definitions of DA, with their discrepancy suggestive of such biases (Supplementary Table 4).

In this regard, an alternative method of dealing with the trade-off between local and global differential abundance that may account for experimental biases would be to allow the user to the specify their estimated ‘true’ proportion of one label and use this as the proportion to test against in GDA. This approach could be extended to allow the user to specify a range of estimated label proportions, with GDA then tested for across this range. We leave these directions for future work.

In any case, Dawnn searches for both LDA and GDA and identifies a higher proportion of DA each than existing state-of-the-art methods, while maintaining a low rate of false discovery (Section 2.3). One design choice that may explain its good performance is that its model inputs are sample labels rather than the label proportions (as used by DA-seq and CNA). This approach allows a flexible model, such as a neural network, to employ the most useful data representation since it can internally convert to a proportion-based representation if that yields greater accuracy (using sample labels as input loses no information, whereas using label proportions does – see Section 12 in Supplementary Information).

Dawnn’s improved performance over Milo might be explained by its cell-level resolution. Its more consistent control of the false discovery rate may be partially due to its use of the Benjamini–Yekutieli procedure, which does not assume independence between the extent of DA expressed by neighbouring cells. We believe Dawnn to be more reliable compared to the only method with similarly high TPRs – CNA – since it better controls the FDR.

Whilst we have shown Dawnn to outperform existing methods following benchmarking procedures established by Dann *et al.* [1], there is a risk that its good performance arises simply because it has been trained to estimate using a training dataset generated in too similar a manner to the benchmarking datasets, thereby biasing the process in its favour. Clearly, an algorithm trained on a dataset similar to the one used for bench-marking may yield good performance as a result of biased training. For this reason, we removed as many assumptions as possible from the training set generation (Section 4.2.2), thus minimising bias towards the benchmarking one. Further, we assessed Dawnn’s performance on a range of single-cell datasets (Table 1) to ensure that its good performance generalises to unseen data. We believe that this our benchmarking dataset^2^ will allow other DA detection methods to be robustly assessed.

Assessing Dawnn’s ability to recover published findings reveals further advantages over existing methods (Section 2.5). Firstly, the range of local log-fold changes between labels for which cells are called as DA is smaller in Dawnn than Milo and DA-seq, despite both Dawnn and Milo attempting to be as conservative as one another (i.e. to stay under an FDR of 10%). Therefore, Dawnn, particularly in its local configuration, identifies more DA populations of interest whilst preserving a low false discovery rate (as seen in Section 2.3). Second, the single-cell resolution of Dawnn and DA-seq allows them to detect more sub-populations exhibiting DA than is possible with the neighbourhood-resolution Milo, likely since neighbourhood-level averaging of fold change estimates leads to signal from small populations being lost. In the liver cirrhosis dataset, Dawnn identified over 2,300 cells as belonging to regions of differential abundance that were missed by DA-seq. Importantly, we observed an increase in the number of such cells identified across a range of cell types (the majority of cell types showed an increase of at least 4%, maximum increase 69%). Detecting differential abundance in rare cell types is crucial as they could be key to understanding some diseases (e.g. ionocytes in COPD and cystic fibrosis [18]). Dawnn-L’s good performance on rare cell types allows it to capture biological insights that are missed by existing methods, as demonstrated for ionocytes and other rare cell types in COPD patients.

We found in Section 2.6 that the runtime of Dawnn is similar to that of Milo, and is slower than CNA but faster than DA-seq. We believe that its slightly larger runtime is justified since it allows the single-cell resolution of the data to be exploited fully. Since it remains possible to run Dawnn in a matter of minutes on a standard laptop, we believe this slowdown to be of little significance in practice.

The pre-trained nature of Dawnn comes at the expense of flexibility in terms of experimental design. Unlike Milo, Dawnn can only be used to assess differential abundance in studies involving two samples or conditions (a limitation shared by DA-seq). Further, Milo can control for batch effects by incorporating covariates related to the experimental design in its statistical model. Dawnn’s pre-trained model again prevents this, although a batch-corrected dimensionality reduction can be used when constructing the KNN graph (in fact, one is used for the growth-accelerated keratinocyte dataset). Similarly, Dawnn’s pre-trained model requires a nearest neighbour graph of 1,000 neighbours for each cell, making the choice of the *K* parameter hard-coded. We chose a large neighbourhood size as this allows differential abundance to be detected at a range of scales (as in DA-seq). Dawnn’s performance may be improved by allowing this parameter to be optimised by the user for their dataset. Despite these decreases in flexibility in terms of experimental design and parameter choice, we believe that Dawnn’s ability to identify DA with higher accuracy than existing methods means that it will prove invaluable in the typical assessment of two samples or experimental conditions. Further, since Dawnn’s training code is open-source^3^, its neural network can be retrained with *K* ≠ 1000 if desired.

When designing Dawnn, we investigated whether one-dimensional convolutional neural networks or graph neural networks [19] could reduce the error when estimating 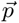 by exploiting local neighbourhood structure. However, we found that this was not the case, at least in our hands. One explanation for this is that this was caused by the lack of structure in the training data, meaning that performance could not be improved by attempting to exploit it. When training neural networks, it is common to use dropout to improve generalisation of the learned model. However, we found it to have a negative effect when training Dawnn (Table 2), again possibly because there is little opportunity to overfit the training data due to its lack of structure. We therefore believe that the negative effect of dropout indicates, as with the poor performance of convolutional and graph neural networks, that we have succeeded in our aim to create training data with little structure and removed benchmarking bias in Dawnn’s favour.

We conclude that Dawnn’s good performance will allow the identification of additional numbers of cells and cell types with DA, widening the analysis of both common and rare cell populations and their biology at single-cell resolution.

## Supplementary information

Supplementary information is contained in the appendix.

## Data availability

Benchmarking datasets, docker images, and other resources needed to reproduce all results in this paper are available online^4^.

## Code availability

Benchmarking code is available online^4^. Code for Dawnn itself is also available^5^.

## 4 Methods

### 4.1 Dawnn

In this section, we use the notation defined in Section 2.2. In contrast to earlier differential abundance detection algorithms, Dawnn has been trained to quantify 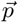 from simulated datasets. We created simulated training data and learned the relationship between each *p* and the labels of the 1,000 nearest neighbours of each cell using a neural network. This trained model can subsequently be applied to any single-cell RNAseq dataset, for example our test datasets in Table 1.

We tested a number of neural network models in order to identify the best-performing architecture. Neural network architectures and the means by which we chose the final one are discussed in the Supplementary Information (Sections 11.1 and 11.2, respectively).

We assessed six neural network architectures, the effect of dropout layers between each hidden layer and the impact of two learning rates. The results of the model selection process are shown in Table 2, in which a lower score indicates better performance during training.

**Table 2:** Results of five-fold cross validation of different neural network architectures, learning rate (LR), and whether dropout. The best-performing algorithm (lowest MSE) is shown in bold. Asterisks indicate incomplete cross validation runs, terminated due to runtime constraints.

| Number of nodes<br>per hidden layer | LR = 0.0001<br>with dropout | LR = 0.0001<br>no dropout | LR = 0.00001<br>with dropout | LR = 0.00001<br>no dropout |
| --- | --- | --- | --- | --- |
| [1000] | 0.001055 | 0.001001 | 0.001855 | 0.001991 |
| [1000,1000] | 0.070437 | 0.000939 | 0.071612 | 0.001094 |
| [250,50,10] | 0.008458 | 0.001107 | 0.007028 | 0.001809 |
| [1000,250,50,10] | 0.006757 | 0.000683 | 0.005723 | 0.001000 |
| [1000,2000,<br>1000,250,10] | 0.006159 | 0.00066 | 0.00787* | 0.000654 |
| [1000,2000,4000,<br>2000,1000,250,10] | 0.00676* | 0.000708 | 0.01076* | **0.000617** |

We also investigated whether other machine learning models – support vector machines [20] and random forests [21] – provided a better basis to estimate 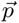 (Table 3). The cross-validated mean squared errors for each model exceed those of the better-performing neural networks, indicating worse performance.

**Table 3:** Results of five-fold cross validation of different statistical models.

| Model | Mean squared error |
| --- | --- |
| Support vector machine | 0.00214 |
| Random forest (100 trees, 50 features) | 0.00346 |
| Random forest (500 trees, 50 features) | 0.00332 |
| Random forest (100 trees, 250 features) | 0.00339 |
| Random forest (500 trees, 250 features) | 0.00324 |

The model selection process identified a neural network with seven hidden layers (containing [1000 2000 4000 2000 1000 250 10] nodes) as having the best performance. This model was trained with a learning rate of 0.00001 and without employing dropout. We then trained this model on the whole training set and incorporated it as the model used by Dawnn.

A more detailed description of neural networks and overview of implementation details of the practical implementations of Dawnn is given in Section 11 in Supplementary Information.

#### 4.1.1 Training set generation

Since Dawnn requires training data to learn to estimate 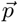, there is a risk that good benchmarking performance may only be indicative of similarity between the training and test datasets. If the same assumptions made during training are also satisfied during benchmarking, then good benchmarking performance may not generalise to real data. It is therefore crucial to minimise bias in the training set in order to learn a generalisable model and enable rigorous, objective, and biologically relevant benchmarking. To this end, we ensured that:

1. There is no global underlying structure to the cell population of the training set;
2. There is only a weak relationship between the labels of neighbouring cells.

If these conditions are satisfied, then Dawnn should learn how to estimate 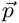 in a generalisable manner since the assumptions used when generating the training dataset are so weak.

To satisfy the first condition, we generated each training instance as an independent 1000-cell dataset, rather than by simulating labels on an larger single-cell dataset (which would have introduced assumptions about the dataset’s structure).

To satisfy the second condition, we constructed each training instance in a stochastic manner, whereby the *p* associated with a cell is generated from the *p* associated with a neighbouring cell by adding a number randomly chosen from a given range, the bounds of which are also randomly chosen for each training instance. The values generated by this process can be thought of as representing the *p*s of cells along a one-dimensional differentiation trajectory. The pseudocode for the training set generation is given in Algorithm 2. The 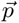 vectors corresponding to the first ten members of the training set are shown in Fig. 7.

##### Algorithm 2: Training set generation.

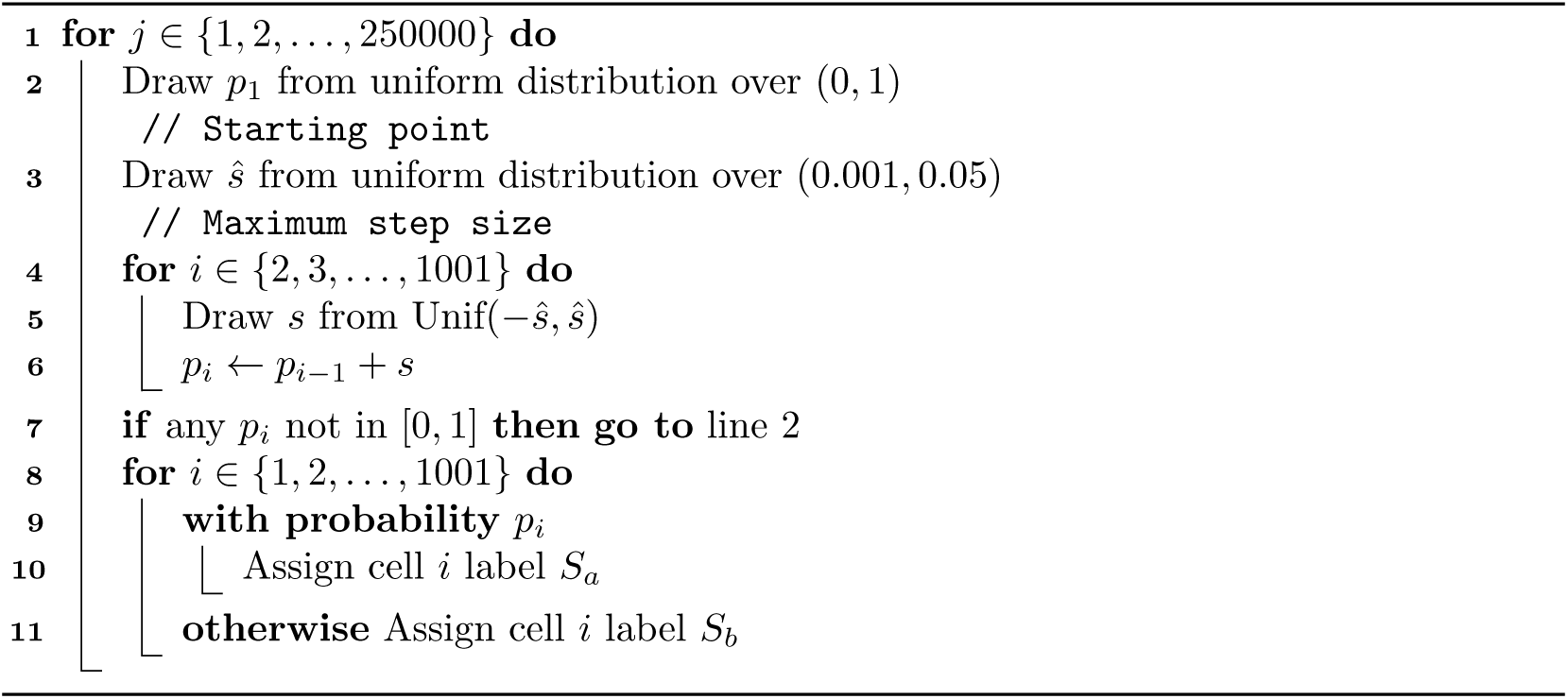

Finally, we tested Dawnn in a wide variety of simulated and real-word datasets (Table 1) with different underlying structures to ensure wide applicability to biological problems.

**Fig. 7:**
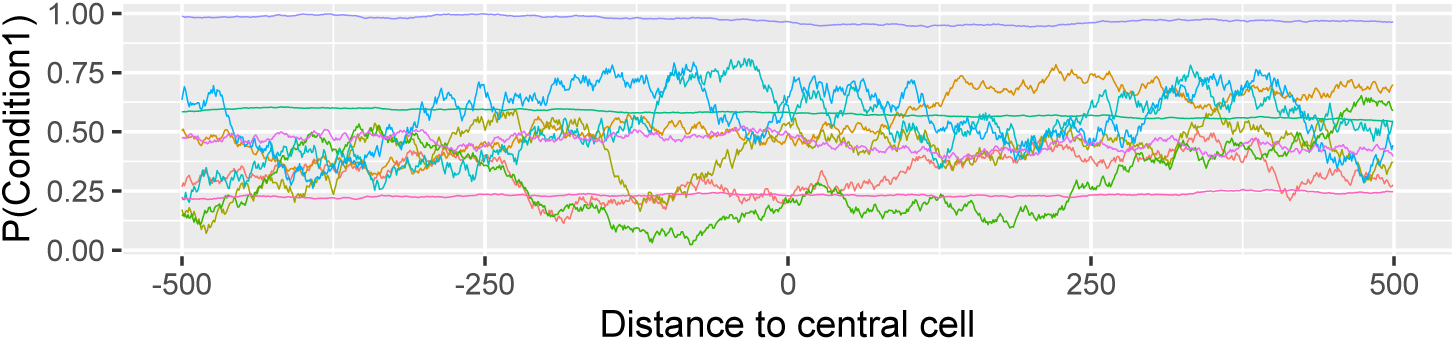
The first ten 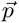 in the training set. The parameters of the random walk dictate the variation in the distribution across each 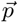.

### 4.2 Benchmarking

#### 4.2.1 Other DA detection methods

The parameter values used for benchmarking existing methods are given in Section 13 in Supplementary Information. Where possible, they were based off those used by Dann *et al.* [1].

##### Milo

Milo [1] creates overlapping neighbourhoods on a *K*-nearest neighbour graph and tests each neighbourhood for differential abundance by fitting a negative binomial generalised linear model to the counts of different labels. It computes p-values using an F-test. Milo does not natively test for differential abundance at single-cell resolution. For the benchmarking in Section 2.3, we follow Dann *et al.* in converting Milo’s neighbourhood level estimates to single-cell resolution by considering “the average outcome in neighbourhoods to which a cell belongs” [1].

##### DA-seq

For each cell, DA-seq [2] computes the proportion of cells with a given label within neighbourhoods of different sizes. It fits a logistic regression model to these proportions and uses this model to estimate the probability that the cell belongs to the given sample (i.e. p in the terminology of Section 2.2). It classifies as DA all cells with a more extreme estimated p than the estimates obtained after shuffling labels.

##### CNA

The statistical framework of CNA [3] requires multiple samples associated with each label. For each cell, it it creates a neighbourhood containing all cells likely to be reached in a random walk starting at that cell and measures the abundance of each sample in the neighbourhood. It then estimates the differential abundance in each neighbourhood as the correlation between the abundance of each sample and their associated labels (i.e. whether there is either a high or low abundance of cells with a specific label). FDRs are estimated by permuting labels to obtain null distributions and neighbourhoods exhibiting DA are identified as those for which the estimated FDR does not exceed a given threshold.

#### 4.2.2 Generating Benchmarking Datasets

We employed a slightly modified version of the benchmarking methodology used by Dann *et al.* when assessing Milo [1], using their code^6^ wherever possible. We followed them in using the mouse gastrulation benchmarking dataset and employed their approach to simulate cells in both discrete clusters and in linear and branching trajectories. We used the same approach as for the mouse gastrulation dataset when assigning simulated labels to the keratinocyte, organoid, and heart datasets.

The benchmarking data was simulated under the same underlying model as that assumed by Dawnn, i.e. that there are two labels (S*_a_* and S*_b_*) and each cell has a probability *p* of being assigned label *S_a_* and labels are sampled according to these probabilities. Different benchmarking instances were simulated by creating distributions of *p* over a dataset by first choosing a cluster (generally corresponding to cell types in the biological datasets) in which the abundance of label *S_a_* is maximally increased and then choosing the maximum *p* (i.e. bias towards *S_a_*) within this cluster. The *p* each cell is calculated proportionally to its distance from the centre of the chosen up-regulated cluster. Each cell is then assigned the label *S_a_* with probability *p* and the label *S_b_* otherwise. We made the same choices as Dann *et al.* [1] when selecting both which clusters to exhibit maximal differential abundance and the maximum *p* in each datasets. To avoid labelling cells with *p* close to 0.5 as ‘not DA’, when, at least under LDA, there is little DA to detect, we set the *p* of cells with *p* ≤ 0.55 to 0.5.

Two ground truth labels were simulated, corresponding to LDA and GDA and following the definitions in Section 5.1 (Supplementary Information) with tolerance *ϵ* = 0.05. Explicitly, for LDA, all cells with *p* outside [0.5 − *ϵ*, 0.5 + *ϵ*] were labelled as exhibiting LDA in the ground truth. For GDA, all cells with *p* outside [*L* − *ϵ*, *L* + *ϵ*] were labelled as exhibiting GDA in the ground truth, where *L* is the global proportion of label *S_a_*.

## Acknowledgements

GTH and SC are funded by the NIHR Great Ormond Street Biomedical Research Centre. We would like to acknowledge that benchmarking in this work was largely based on that from Dann *et al.* [1], whose clear methodology and code we appreciated enormously. We would also like to acknowledge insightful discussion with the authors of CNA [3] around the subtleties in the definition of differential abundance and its application in simulation. Ultimately, these discussions led to our creation of the more nuanced definitions of local and global differential abundance and subsequent more rigorous benchmarking. We thank Constantin Ahlmann-Eltze for useful discussions.

## SUPPLEMENTARY INFORMATION

### 5 Comparison between local and global differential abundance

In this section, we derive results about the relationship between local and global differential abundance (LDA and GDA, respectively). In particular, we show the impact of uneven numbers of labels on whether a cell population is considered to exhibit LDA or GDA. We also outline which definitions are used in the literature.

#### 5.1 Definitions

In the following, we assume that the dataset comprises exactly two labels, *L*_1_ and *L*_2_, and that the number of cells with each label is *n*_1_ and *n*_2_, respectively. We assume also that each cell *c_i_* has an underlying probability *p_i_* of being assigned label *L*_1_ (and hence a probability of 1 *− p_i_* of being assigned label *L*_2_). We assume that the observed labels are an instantiation of this generative distribution. We incorporate the notion of statistical significance by introducing an interval of *ϵ* around the threshold on *p_i_* for differential abundance outside of which cells are not considered to exhibit DA.

In this setting, we can now define local and global differential abundance.

**Definition 5.1** (*ϵ*-local differential abundance). For a tolerance 0 ≤ *ϵ* ≤ 0.5, the set of cells exhibiting ϵ-*local differential abundance* (*ϵ*-LDA) is

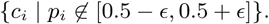

**Definition 5.2** (*ϵ*-global differential abundance). For a tolerance 0 ≤ *ϵ* ≤ *n*_1_/(*n*_1_ +*n*_2_), the set of cells exhibiting ϵ-*global differential abundance* (*ϵ*-GDA) is

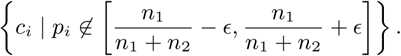

Note that if *n*_1_ = *n*_2_ (i.e. when there is no class imbalance), then the definitions of *ϵ*-GDA and *ϵ*-LDA are equivalent: *ϵ*-LDA is a special case of *ϵ*-GDA where there are the same number of cells with each label.

In this work, the exact value of *ϵ* is often unimportant and thus can be omitted in favour of simply calling the two definitions ‘LDA’ and ‘GDA’.

#### 5.2 Class imbalance

A dataset with labelled cells exhibits a class imbalance if there are more cells with one label than with the other. In our analysis of the relationship between LDA and GDA, we use two notions of class imbalance: global and local.

**Definition 5.3** (Global class imbalance). A dataset exhibits a *global class imbalance* if and only if there are unequal numbers of cells with each label.

**Definition 5.4** (Local class imbalance). A dataset exhibits a *local class imbalance* if and only if there are neighbourhoods of similar cells (e.g. cell types) with unequal numbers of cells with each label.

Note that if there exists a subset of cells that exhibit LDA, then the dataset exhibits a local class imbalance. Note also that the existence of local class imbalances does not imply the existence of a global one: two equally sized local class imbalances which favour different labels may cancel each other out at the global level.

#### 5.3 Effect of class imbalance on local and global DA

We now investigate the effect of class imbalance on which cell populations exhibit LDA and GDA. For simplicity, we do so in a setting with three clusters of cells. In the tables below, we display the number of cells in each cluster, the proportion of cells with label *L*_1_ in each cluster, and finally the differences between this proportion and the threshold proportion at which a cell population is considered to exhibit DA under LDA and GDA (i.e. 0.5 in LDA and the global proportion of *L*_1_ in GDA). These ‘scores’ signify the strength of differential abundance under each definition. We assume that *ϵ* = 0.05 (as defined in Section 5.1), and therefore that any population of cells with a proportion of *L*_1_ labels that differs from this threshold by more than 0.05 are considered to exhibit DA.

##### 5.3.1 Local class imbalance

In this setting, Clusters 1 and 2 have equal proportions of labels whilst Cluster 3 has a higher proportion of *L*_1_. When the number of cells in Cluster 3 is similar to that in the other two, the same clusters are considered to exhibit DA under both LDA and GDA (Supplementary Table 1).

**Supplementary Table 1:** LDA and GDA classify the same cell populations as exhibiting differential abundance when the cluster of cells exhibiting local class imbalance is a similar size to the other clusters.

| Cluster | Num $L_1$ | Num $L_2$ | Prop $L_1$ | LDA score | LDA | GDA score | GDA |
| --- | --- | --- | --- | --- | --- | --- | --- |
| 1 | 50 | 50 | 0.50 | 0.00 | No | 0.00 | No |
| 2 | 50 | 50 | 0.50 | 0.00 | No | 0.00 | No |
| 3 | 6 | 4 | 0.60 | 0.10 | Yes | 0.10 | Yes |
| 1 | 50 | 50 | 0.50 | 0.00 | No | -0.03 | No |
| 2 | 50 | 50 | 0.50 | 0.00 | No | -0.03 | No |
| 3 | 60 | 40 | 0.60 | 0.10 | Yes | 0.07 | Yes |

However, when the size of the Cluster 3 increases (with 600 *L*_1_ cells and 400 *L*_2_ cells), LDA and GDA classify different cell populations as exhibiting DA (Supplementary Table 2). Under LDA, Cluster 3 exhibits DA (with *L*_1_ differentially abundant), whilst Clusters 1 and 2 do not. Under GDA, on the other hand, Clusters 1 and 2 exhibit DA (with *L*_2_ differentially abundant) whilst Cluster 3 does not.

**Supplementary Table 2:**
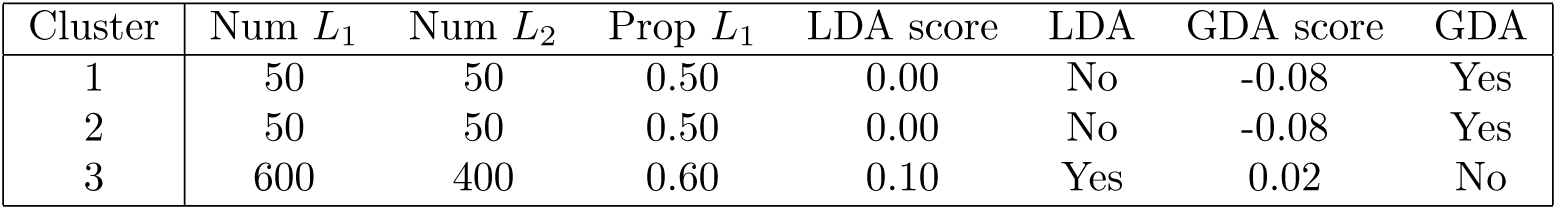
LDA and GDA classify different cell populations as exhibiting differential abundance when the cluster of cells exhibiting local class imbalance is much larger than other clusters.

Supplementary Fig. 1 shows the effect of increasing the size of Cluster 3, from ten to 1,000 cells whilst the sizes of the other two clusters remains 100 cells. For each cluster, we assume a 60/40 split in favour of *L*_1_. For each dataset, we plot the LDA and GDA ‘scores’, as calculated in the above tables. We can see that, once the number of cells in Cluster 3 is approximately 160, the GDA score falls below 0.05, whilst the score for LDA stays around 0.1. Therefore, if there are more than 160 cells in Cluster 3, then, for *ϵ* = 0.05, Cluster 3 is deemed to exhibit differential abundance under LDA but not under GDA. Therefore, for large local class imbalances, LDA and GDA classify different cell populations as exhibiting DA.

**Supplementary Fig. 1:**
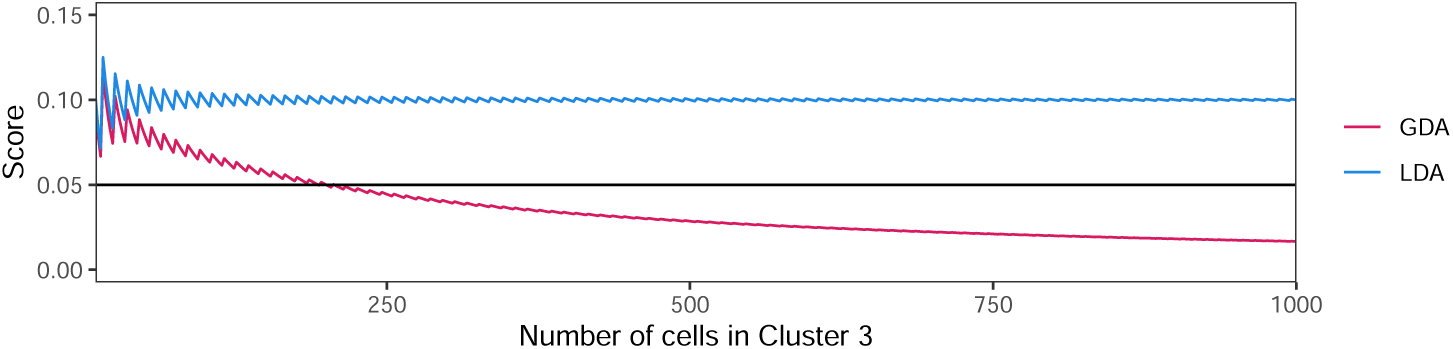
LDA and GDA classify different cell populations as exhibiting differential abundance when the cluster exhibiting a local class imbalance is large enough. Given 100 cells in Clusters 1 and 2, when Cluster 3 contains more than 160 cells, it exhibits LDA but not GDA (using a minimum score threshold of 0.05, indicated by the horizontal line).

##### 5.3.2 Global class imbalance

We now consider a global class imbalance, where each cluster has twice as many *L*_1_ cells as *L*_2_. Here, each cluster exhibits DA under LDA but none do so under GDA (Supplementary Table 3).

**Supplementary Table 3:** LDA and GDA classify different cell populations as exhibiting differential abundance under global class imbalances.

| Cluster | Num $L_1$ | Num $L_2$ | Prop $L_1$ | LDA score | LDA | GDA score | GDA |
| --- | --- | --- | --- | --- | --- | --- | --- |
| 1 | 100 | 50 | 0.67 | 0.17 | Yes | 0.00 | No |
| 2 | 100 | 50 | 0.67 | 0.17 | Yes | 0.00 | No |
| 3 | 200 | 100 | 0.67 | 0.17 | Yes | 0.00 | No |

#### 5.4 Complementary relationship between LDA and GDA

In some settings, as shown above, both LDA and GDA classify the same cell populations as exhibiting DA. However, in other contexts, these cell populations differ. As seen in Section 1, GDA can sometimes lead to false positives whilst LDA identifies the perturbed cell population of interest to the user. However, if there are large global class imbalances spread evenly across many neighbourhoods, then cell populations may exhibit LDA even if the expression within each cell type is identical between samples. This can occur, for instance, if twice as many cells have been sequenced from one sample than the other. A number of experimental biases can cause this situation, such as differences in sampling, processing, or preservation methods. Under GDA, on the other hand, no cell populations would be considered to exhibit DA in this setting. It is not possible to handle this issue computationally, e.g. through downsampling, since it is unclear how to identify and mitigate the experimental bias responsible. Nevertheless, this impact of experimental bias means that, by searching for both LDA and GDA concurrently, the user can diagnose experimental biases according to Supplementary Table 4. We have designed Dawnn to perform well in both the detection of LDA and GDA, allowing the user to investigate both.

**Supplementary Table 4:**
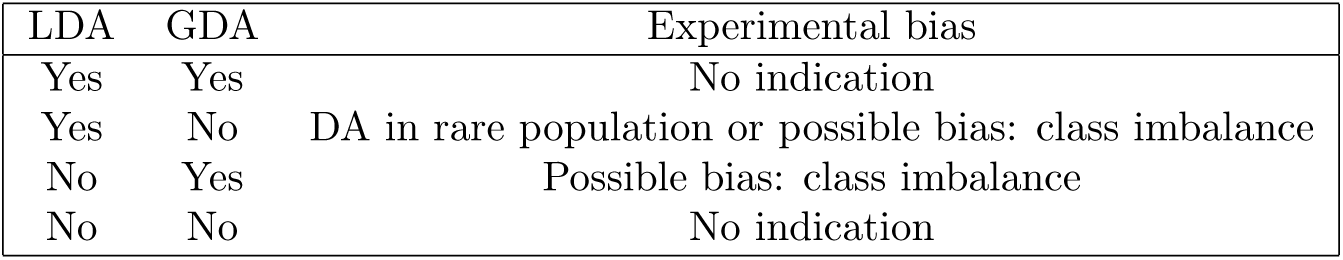
Diagnosis of experimental bias by searching for LDA and GDA concurrently.

#### 5.5 Local and global DA detection in literature

Let *X* := {*x*_1_, …, *x_n_*} ∈ ℝ*^m×n^* be the gene expression matrix for *n* cells where the expression of *m* genes has been measured and let gene *x_i_* have label *y_i_*.

##### 5.5.1 DA-seq

DA-seq aims to identify global differential abundance. Following the write-up in [2], it defines two probability density functions, *f*_1_ and *f*_2_ over ℝ*^m^* as

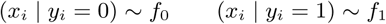

and aims to find regions of *R^m^* where one function is significantly larger than the other. It assumes that each cell is assigned label *L*_1_ with probability *ρ* and it is otherwise assigned label *L*_2_. DA-seq computes a statistic *d*(*x*) for the differential abundance in the neighbourhood of a cell *x* with label *y*:

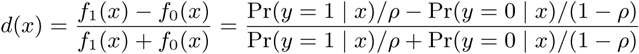

The presence of the Bernoulli parameter *ρ* means that this statistic is calculated in terms of the global ratio between the two labels, making DA-seq a global method.

##### 5.5.2 Milo

Milo creates a set of overlapping neighbourhoods of cells. It then computes a matrix *M* that contains, for each neighbourhood, the number of cells belonging to each sample. It identifies differentially abundant neighbourhoods using the differential expression testing framework from edgeR [22]. Effectively, Milo treats each neighbourhood in *M* as a ‘gene’ and tests which are ‘differentially expressed’ (in our context, differentially abundant) in one set of samples compared to another.

The edgeR statistical framework employs the trimmed method of means (TMM) to normalize cell counts in *M* (‘library sizes’ in the RNA-seq literature of TMM). Briefly, in TMM, counts are multiplied by a *normalisation factor* computed from the average log-fold change in neighbourhoods without an extreme log-fold change between samples and where neither sample is extremely highly abundant. When calculating their average, each log-fold change is weighted by the reciprocal of the variance of the sample counts within its neighbourhood.

If Milo normalised cell counts without TMM (by scaling down counts from the more abundant sample until they have the same total as the less abundant, i.e. running TMM without trimming), then it would behave similarly to DA-seq since it compensates for a global class imbalance. With TMM, Milo still behaves similarly to global methods since it compensates for class imbalance, albeit with a small change in how it calculates the factor by which it scales down the counts of cells from more abundant samples. Regardless of the amount of trimming employed by TMM, Milo does not run as a local method. We therefore claim that Milo operates in between local and global settings.

##### 5.5.3 CNA

CNA [3] first creates a neighbourhood around each cell, where the probability that cell *A* belongs to the neighbourhood of cells *B* is proportional to the probability that *A* would be reached from a random walk starting at *B* (according to some definition of transition probabilities, e.g. a UMAP graph). It then computes the neighbourhood abundance matrix (NAM) in two steps. First, a matrix *R* is constructed whose *n, m*-th entry is the expected number of cells from the *n*-th sample that would arrive in the *m*-th neighbourhood following a random walk. Each row (corresponding to a sample) of *R* is then normalized to sum to 1, yielding the relative abundance of each cell’s neighbourhood in each sample: the NAM.

CNA computes a ‘smoothed correlation’ between the columns of the NAM (i.e. cell neighbourhoods) and the vector *y* of sample-level attributes, e.g. case-control labels. It then shuffles *y* to create a null distribution and test statistical significance of differential abundance within each cell’s neighbourhood.

CNA cannot easily be classified as either a local or global method since the label shuffling for the null distribution takes places at the sample rather than cell level, meaning that class imbalances are hidden in these sample-level statistics. As with Milo, some edge cases can be distinguished. For example, at one extreme, if each cell were assigned to its own sample then CNA would behave as a global method as shuffling would be at the cell level.

### 6 Benchmarking for global DA

This section includes benchmarking results when searching for global DA. Supplementary Fig. 2 contains results for the fully simulated datasets and Supplementary Fig. 3 contains results for the experimental datasets.

**Supplementary Fig. 2:**
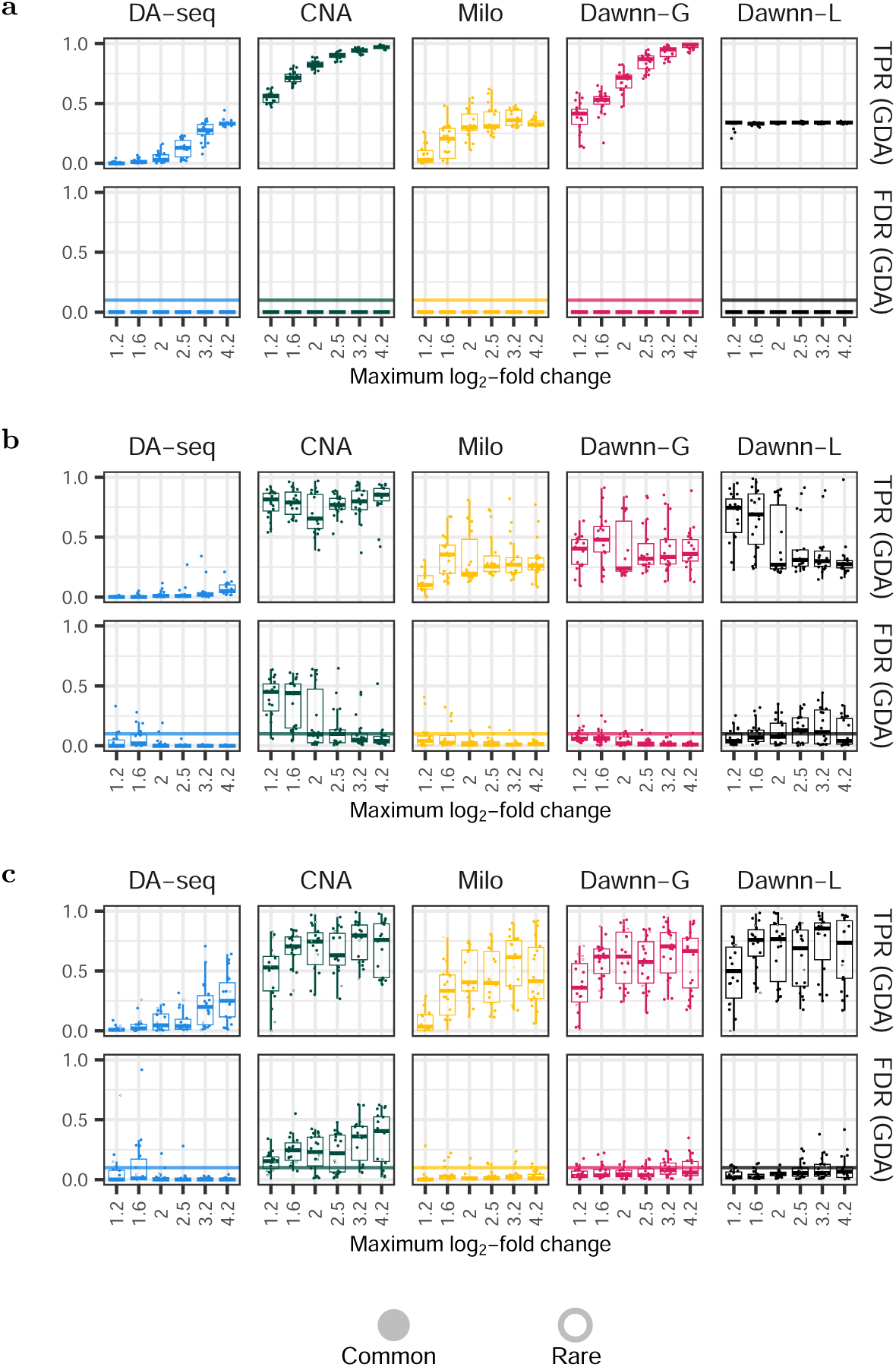
True positive and false discovery rates on simulated datasets for LDA. **a** discrete clusters; **b** linear trajectories; **c** branching trajectories. Filled circles correspond to common cell types (>5% abundance); white circles to rare (*≤*5%). Dawnn-G consistently identifies the (sometimes joint-) largest proportion of cell populations exhibiting GDA whilst maintaining an FDR generally below the 10% threshold (far lower than CNA for all datasets except discrete clusters).

**Supplementary Fig. 3:**
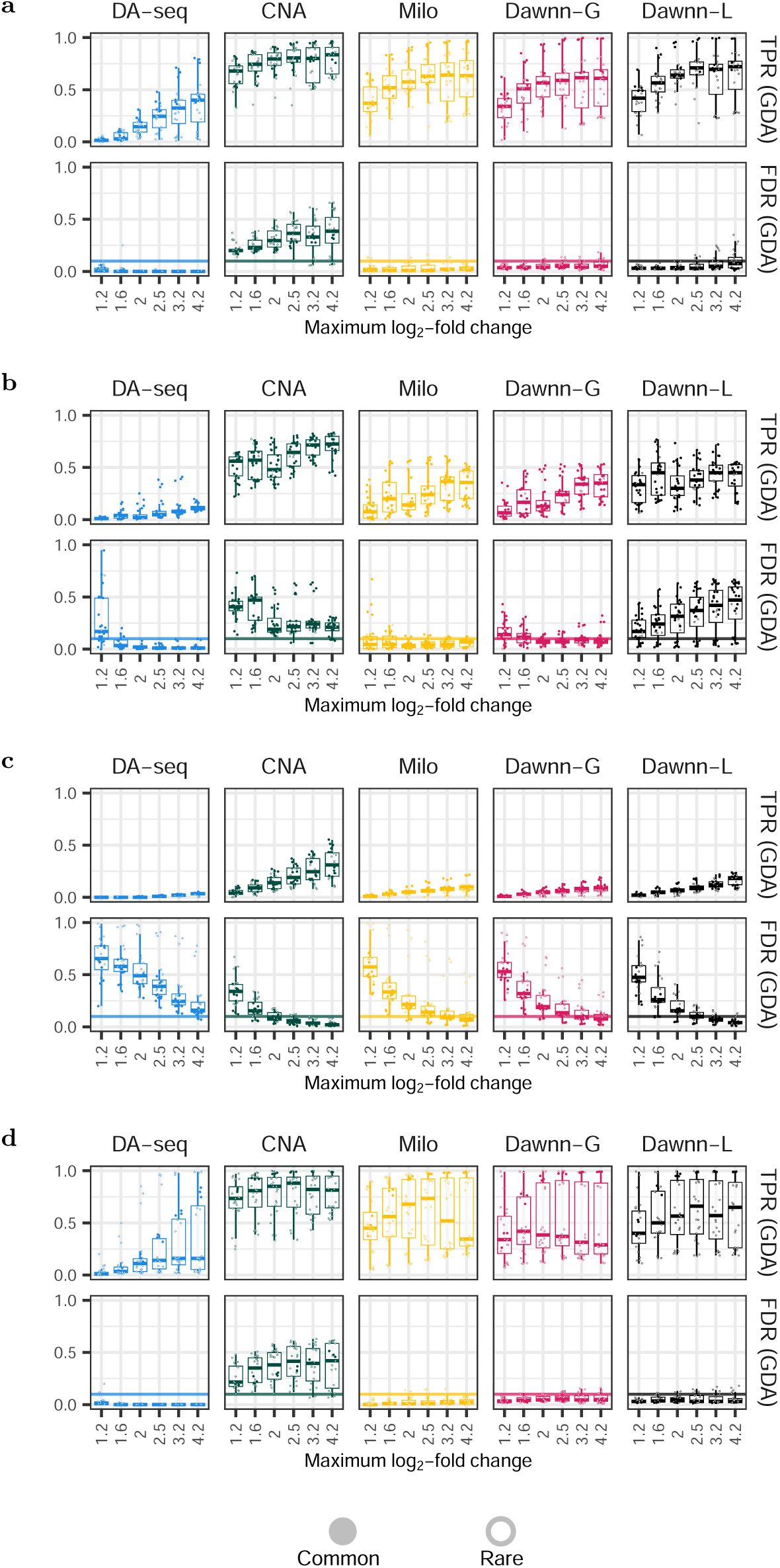
True positive and false discovery rates on simulated datasets for GDA. **a** mouse embryo; **b** keratinocytes; **c** organoids; **d** heart. Filled circles correspond to common cell types (>5% abundance); white circles to rare (*≤*5%). Dawnn-G is the method with highest TPR that controls its FDR below 10%.

### 7 Statistical analyses

Supplementary Fig. 4 aggregates benchmarking runs across all runs and tests for statistically significant differences between methods. Results were compared using a paired Wilcoxon signed-rank test with the Bonferroni correction employed to control for multiple comparisons.

**Supplementary Fig. 4:**
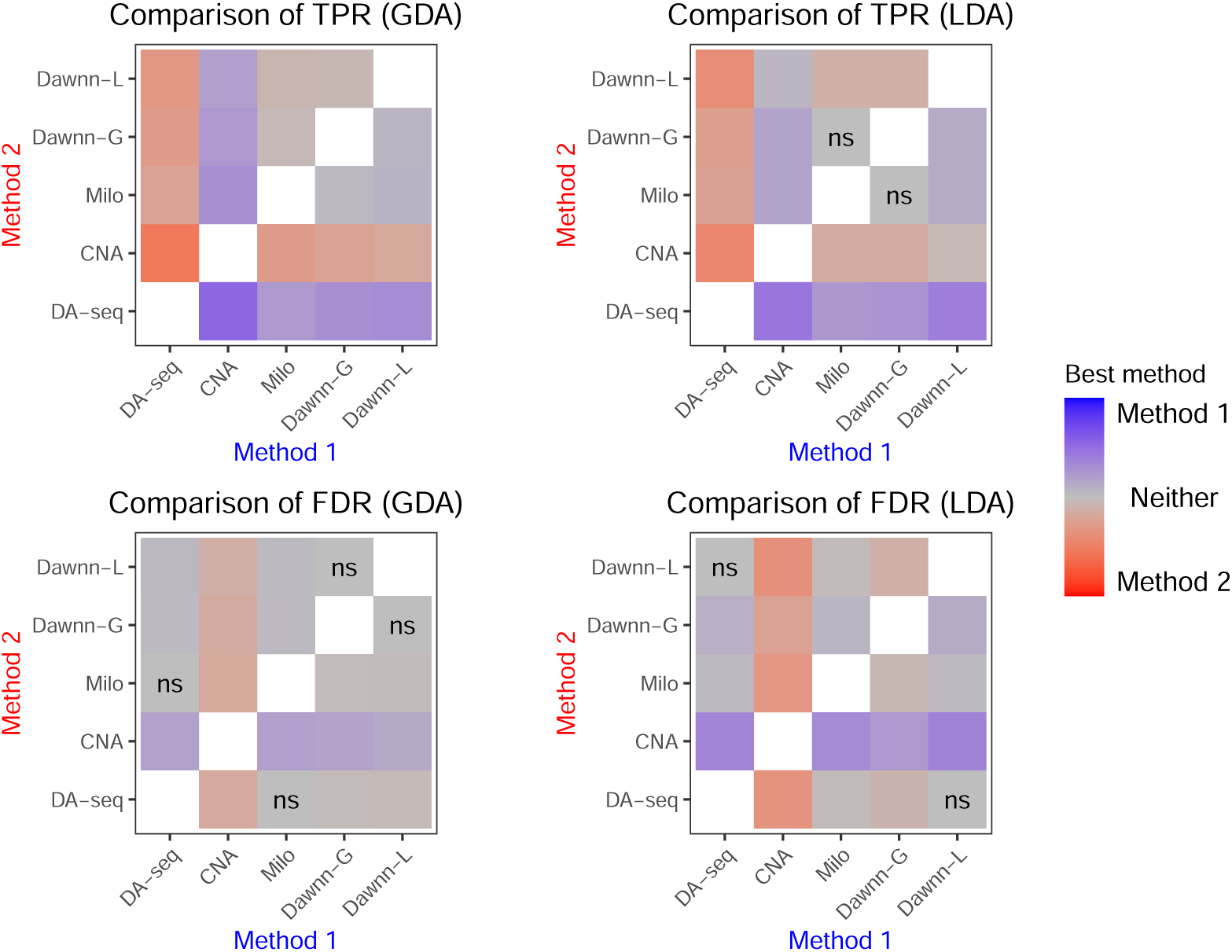
Statistical comparisons of true positive and false discovery rates of DA-seq, Milo, CNA, Dawnn-G, and Dawnn-L across all datasets. Colour corresponds to the estimate of the median of the difference of a sample of TPR or FDR from values for Method 1 and Method 2: darker blue indicates that Method 1 outperformed Method 2, i.e. TPR (FDR) was higher (lower) for Method 1. Darker red indicates the opposite, i.e. Method 2 outperformed Method 1. “ns” indicates no significant difference found.

### 8 Influence of rarity of DA cell population

This section includes benchmarking results both split by rarity of the cell type driving the differential abundance (Supplementary Fig. 5) and with the TPR and FDR plotted in terms of the abundance of the DA-driving cell type (Supplementary Figs. 6 and 7).

**Supplementary Fig. 5:**
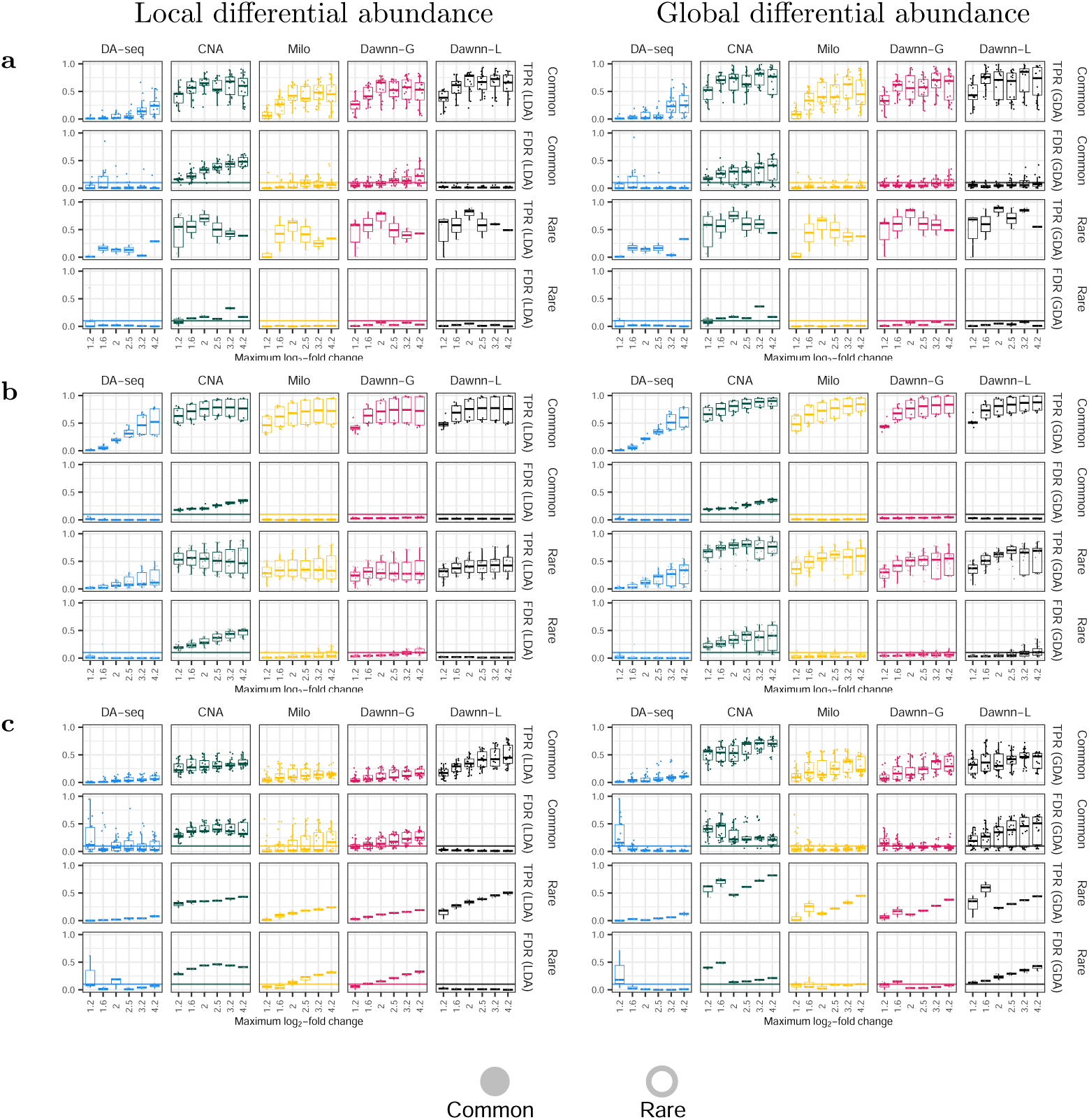
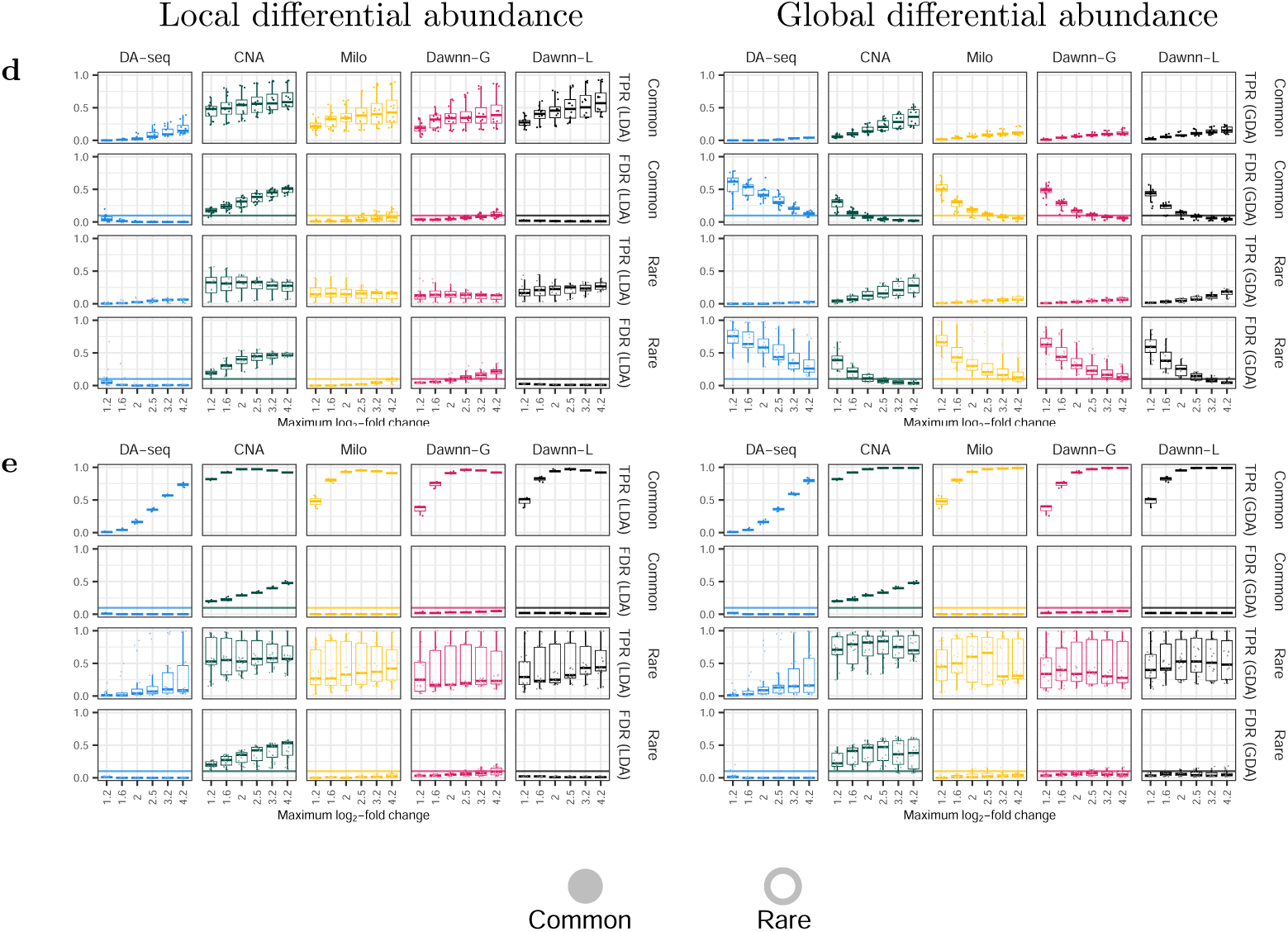
True positive and false discovery rates of DA-seq, CNA, Milo, and both Dawnn approaches for experimental datasets for detecting LDA (left) and GDA (right), split by rarity of differentially abundant cell type. **a** branching trajectories; **b** mouse embryo; **c** keratinocytes; **d** organoids; **e** heart.

**Supplementary Fig. 6:**
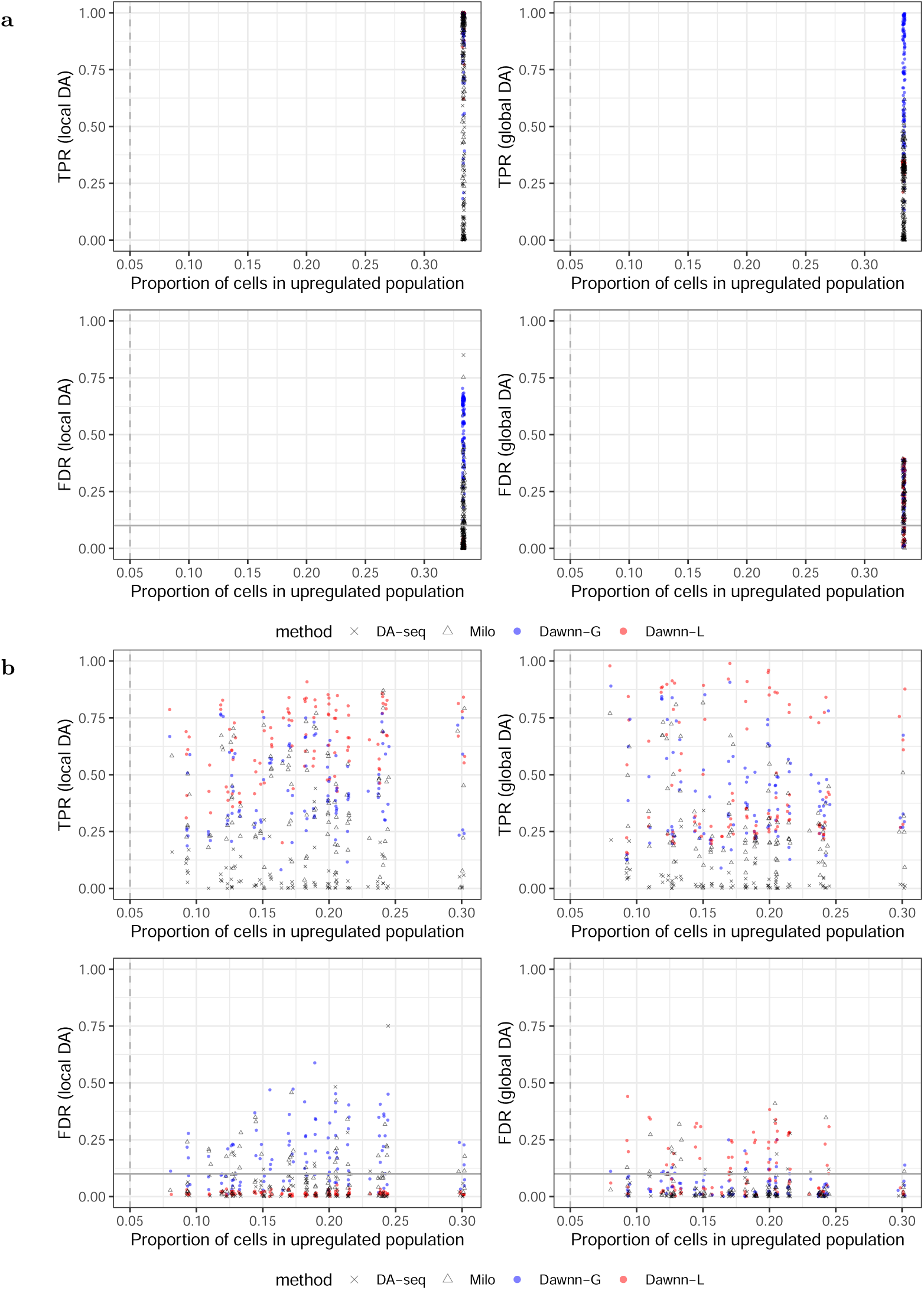
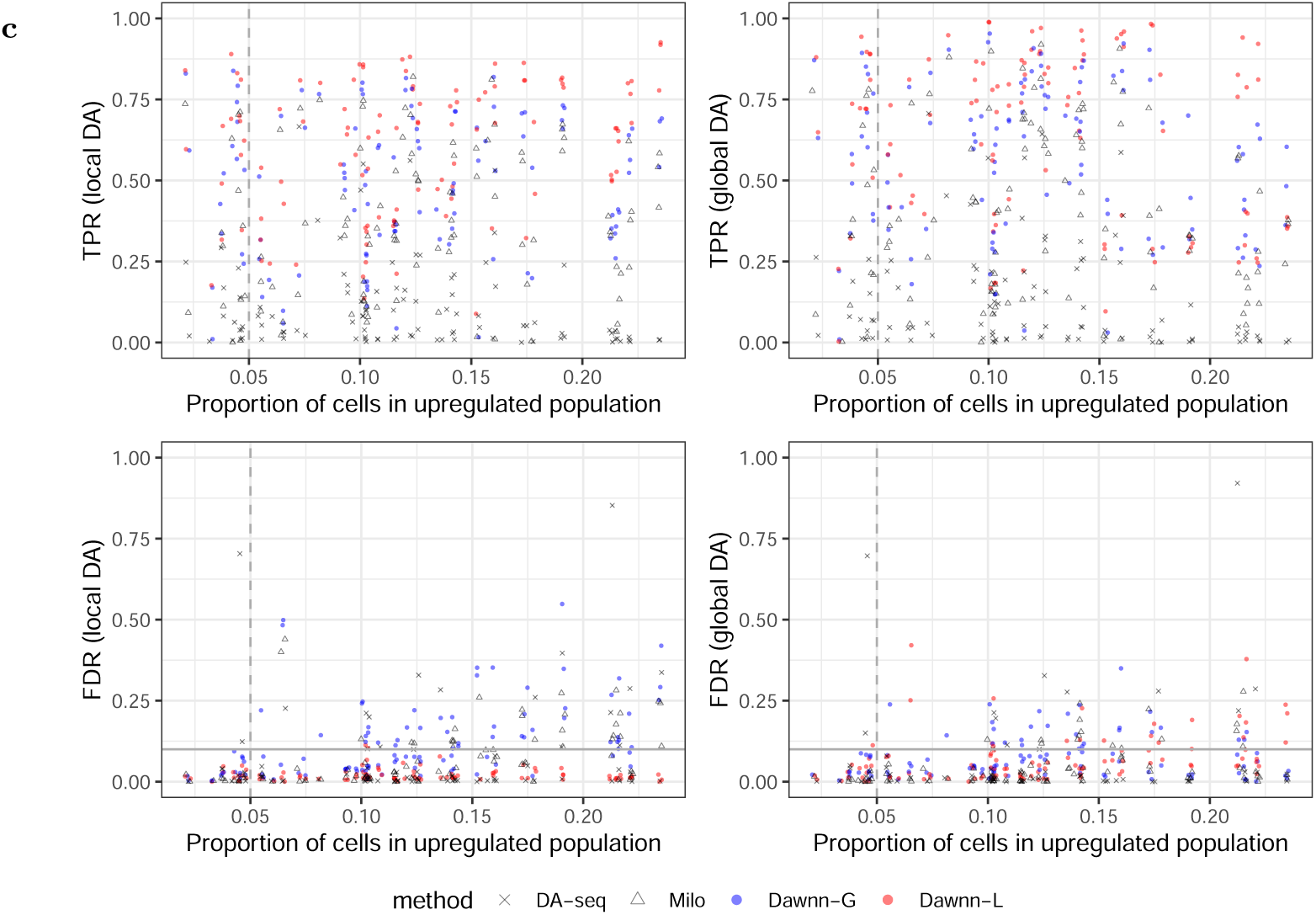
True positive and false discovery rates for simulated datasets for detecting LDA (left) and GDA (right) in relation to rarity of differentially abundant cell type. **a** discrete clusters; **b** linear trajectories; **c** branching trajectories.

**Supplementary Fig. 7:**
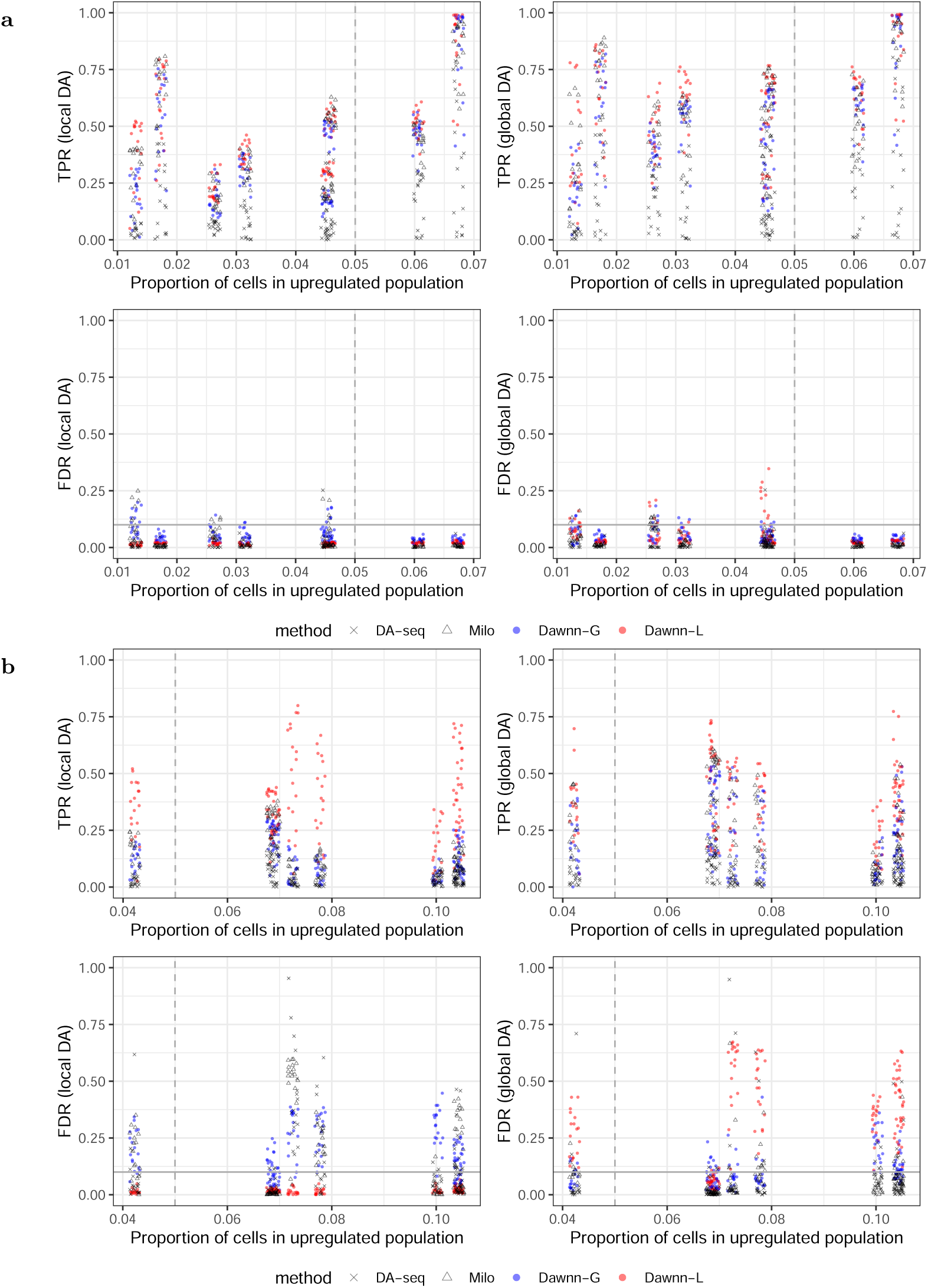
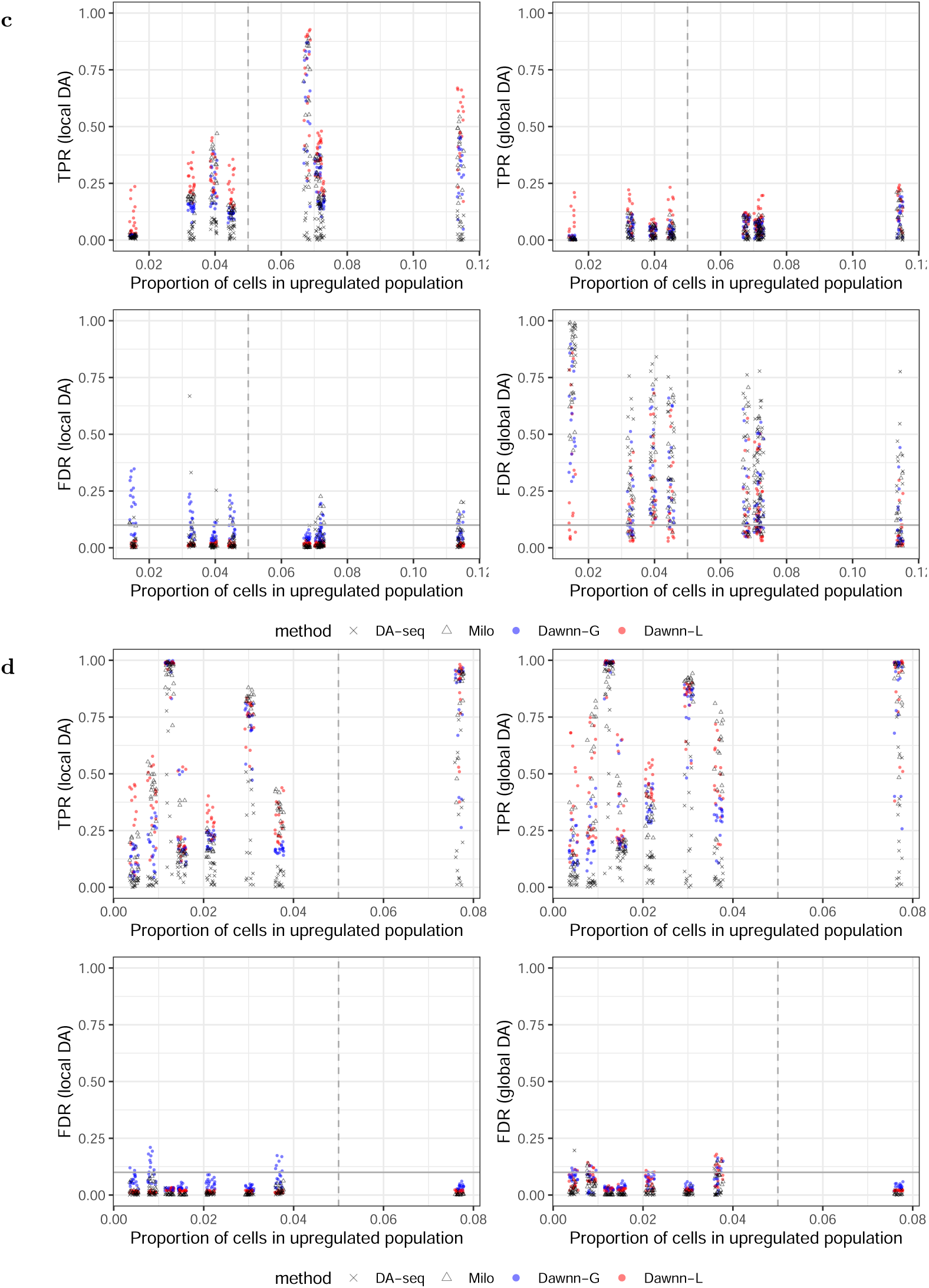
True positive and false discovery rates for experimental datasets for detecting GDA (left) and LDA (right) in relation to rarity of differentially abundant cell type. **a** mouse embryo; **b** keratinocytes; **c** organoids; **d** heart.

### 9 Liver cirrhosis – Extended Results

Supplementary Fig. 8 and Supplementary Table 5 show the discovery rates of DA-seq and Dawnn.

**Supplementary Fig. 8:**
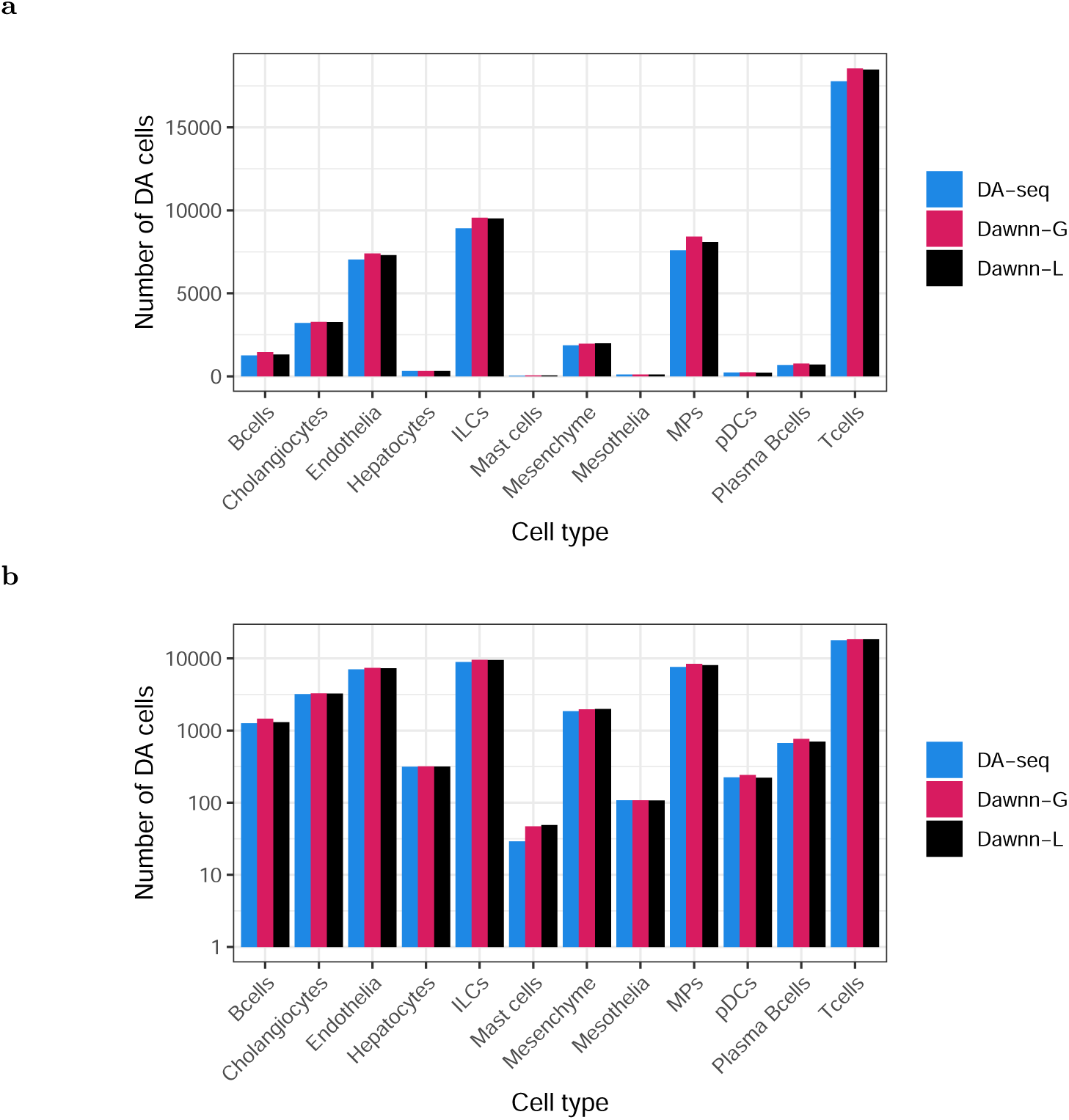
Number of cells in the cirrhotic liver dataset estimated to be in regions of differential abundance by each method, grouped by cell type. For each cell type, both versions of Dawnn estimate at least as many cells as being in DA regions as DA-seq. **a** and **b** use linear and logarithmic y-axes, respectively.

**Supplementary Table 5:** Number of cells estimated to be in regions of DA by each cell-resolution method except CNA (excluded due to high FDR). Percentages are change over number of cells found by DA-seq.

|  | DA-seq | Dawnn-L | Dawnn-G |
| --- | --- | --- | --- |
| Bcells | 1,260 | 1,309 (4%) | 1,458 (16%) |
| Cholangiocytes | 3,205 | 3,266 (2%) | 3,273 (2%) |
| Endothelia | 7,037 | 7,299 (4%) | 7,394 (5%) |
| Hepatocytes | 315 | 317 (1%) | 318 (1%) |
| ILCs | 8,910 | 9,509 (7%) | 9,547 (7%) |
| Mast cells | 29 | 49 (69%) | 47 (62%) |
| Mesenchyme | 1,858 | 1,988 (7%) | 1,966 (6%) |
| Mesothelia | 108 | 107 (-1%) | 108 (0%) |
| MPs | 7,590 | 8,082 (6%) | 8,415 (11%) |
| pDCs | 224 | 222 (-1%) | 242 (8%) |
| Plasma Bcells | 670 | 702 (5%) | 766 (14%) |
| Tcells | 17,769 | 18,481 (4%) | 18,547 (4%) |

### 10 Ionocytes and other rare cell types in COPD – Extended Results

Supplementary Fig. 9 shows results from Section 2.5.2 extended to include all cell types. Supplementary Fig. 10 and Supplementary Table 6 show the discovery rates of DA-seq and Dawnn.

**Supplementary Fig. 9:**
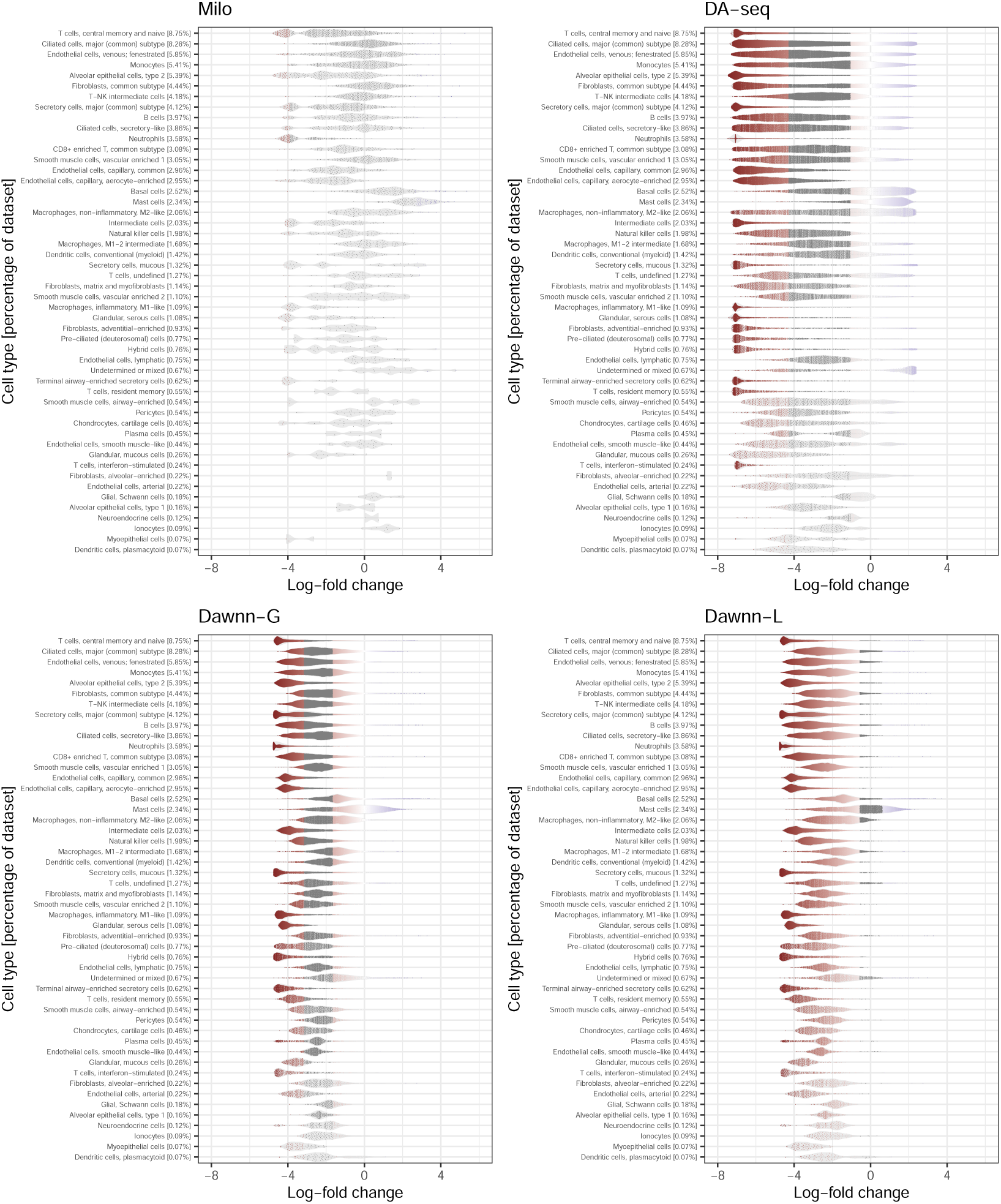
Log-fold changes estimated by Milo, DA-seq, Dawnn-G, and Dawnn-L on the COPD dataset of Rustam *et al.* [5]. Positive log-fold changes correspond to increased abundance in diseased samples; negative to increased abundance in healthy.

**Supplementary Fig. 10:**
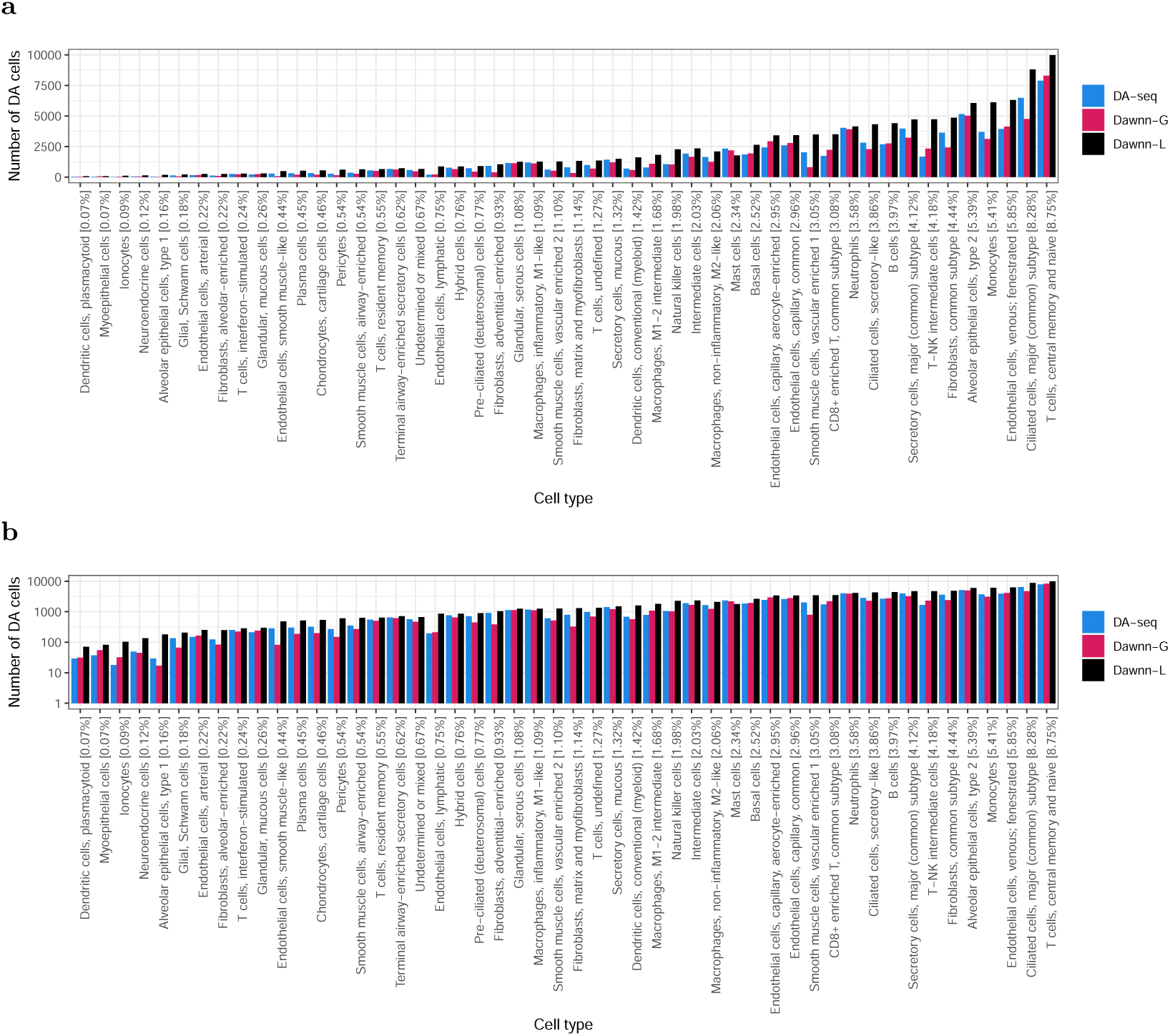
Number of cells in the COPD dataset estimated to be in regions of differential abundance by each method, grouped by cell type. For all cell types except one, Dawnn-L estimates at least as many cells as being in DA regions as DA-seq. **a, b** use linear and logarithmic y-axes, respectively.

**Supplementary Table 6:** Number of cells estimated to be in regions of DA by each cell-resolution method except CNA (excluded due to high FDR). Percentages are change over number of cells found by DA-seq.

|  | DA-seq | Dawnn-L | Dawnn-G |
| --- | --- | --- | --- |
| Dendritic cells, plasmacytoid [0.07%] | 29 | 71 (145%) | 31 (7%) |
| Myoepithelial cells [0.07%] | 37 | 82 (122%) | 55 (49%) |
| Ionocytes [0.09%] | 18 | 103 (472%) | 32 (78%) |
| Neuroendocrine cells [0.12%] | 49 | 135 (176%) | 45 (-8%) |
| Alveolar epithelial cells, type 1 [0.16%] | 29 | 179 (517%) | 17 (-41%) |
| Glial, Schwann cells [0.18%] | 135 | 206 (53%) | 66 (-51%) |
| Endothelial cells, arterial [0.22%] | 149 | 251 (68%) | 165 (11%) |
| Fibroblasts, alveolar-enriched [0.22%] | 124 | 250 (102%) | 84 (-32%) |
| T cells, interferon-stimulated [0.24%] | 252 | 286 (13%) | 223 (-12%) |
| Glandular, mucous cells [0.26%] | 212 | 295 (39%) | 238 (12%) |
| Endothelial cells, smooth muscle-like [0.44%] | 285 | 482 (69%) | 82 (-71%) |
| Plasma cells [0.45%] | 302 | 516 (71%) | 188 (-38%) |
| Chondrocytes, cartilage cells [0.46%] | 324 | 536 (65%) | 199 (-39%) |
| Pericytes [0.54%] | 269 | 607 (126%) | 149 (-45%) |
| Smooth muscle cells, airway-enriched [0.54%] | 356 | 631 (77%) | 269 (-24%) |
| T cells, resident memory [0.55%] | 549 | 638 (16%) | 500 (-9%) |
| Terminal airway-enriched secretory cells [0.62%] | 653 | 715 (9%) | 615 (-6%) |
| Undetermined or mixed [0.67%] | 568 | 664 (17%) | 470 (-17%) |
| Endothelial cells, lymphatic [0.75%] | 195 | 860 (341%) | 213 (9%) |
| Hybrid cells [0.76%] | 761 | 864 (14%) | 646 (-15%) |
| Pre-ciliated (deuterosomal) cells [0.77%] | 715 | 899 (26%) | 447 (-37%) |
| Fibroblasts, adventitial-enriched [0.93%] | 916 | 1,044 (14%) | 383 (-58%) |
| Glandular, serous cells [1.08%] | 1,136 | 1,249 (10%) | 1,134 (0%) |
| Macrophages, inflammatory, M1-like [1.09%] | 1,181 | 1,262 (7%) | 1,109 (-6%) |
| Smooth muscle cells, vascular enriched 2 [1.10%] | 610 | 1,269 (108%) | 516 (-15%) |
| Fibroblasts, matrix and myofibroblasts [1.14%] | 792 | 1,313 (66%) | 325 (-59%) |
| T cells, undefined [1.27%] | 978 | 1,347 (38%) | 694 (-29%) |
| Secretory cells, mucous [1.32%] | 1,418 | 1,497 (6%) | 1,201 (-15%) |
| Dendritic cells, conventional (myeloid) [1.42%] | 690 | 1,604 (132%) | 567 (-18%) |
| Macrophages, M1-2 intermediate [1.68%] | 781 | 1,808 (131%) | 1,085 (39%) |
| Natural killer cells [1.98%] | 1,059 | 2,268 (114%) | 1,028 (-3%) |
| Intermediate cells [2.03%] | 1,917 | 2,334 (22%) | 1,661 (-13%) |
| Macrophages, non-inflammatory, M2-like [2.06%] | 1,638 | 2,097 (28%) | 1,244 (-24%) |
| Mast cells [2.34%] | 2,326 | 1,778 (-24%) | 2,185 (-6%) |
| Basal cells [2.52%] | 1,849 | 2,629 (42%) | 1,931 (4%) |
| Endothelial cells, capillary, aerocyte-enriched [2.95%] | 2,419 | 3,413 (41%) | 2,928 (21%) |
| Endothelial cells, capillary, common [2.96%] | 2,596 | 3,419 (32%) | 2,781 (7%) |
| Smooth muscle cells, vascular enriched 1 [3.05%] | 2,020 | 3,481 (72%) | 800 (-60%) |
| CD8+ enriched T, common subtype [3.08%] | 1,729 | 3,496 (102%) | 2,222 (29%) |
| Neutrophils [3.58%] | 4,026 | 4,139 (3%) | 3,902 (-3%) |
| Ciliated cells, secretory-like [3.86%] | 2,811 | 4,317 (54%) | 2,280 (-19%) |
| B cells [3.97%] | 2,680 | 4,404 (64%) | 2,743 (2%) |
| Secretory cells, major (common) subtype [4.12%] | 3,964 | 4,703 (19%) | 3,218 (-19%) |
| T-NK intermediate cells [4.18%] | 1,669 | 4,727 (183%) | 2,323 (39%) |
| Fibroblasts, common subtype [4.44%] | 3,621 | 4,861 (34%) | 2,413 (-33%) |
| Alveolar epithelial cells, type 2 [5.39%] | 5,143 | 6,053 (18%) | 5,010 (-3%) |
| Monocytes [5.41%] | 3,690 | 6,107 (66%) | 3,109 (-16%) |
| Endothelial cells, venous; fenestrated [5.85%] | 3,932 | 6,304 (60%) | 4,121 (5%) |
| Ciliated cells, major (common) subtype [8.28%] | 6,471 | 8,807 (36%) | 4,747 (-27%) |
| T cells, central memory and naive [8.75%] | 7,883 | 9,977 (27%) | 8,287 (5%) |

### 11 Machine learning details

This section provides more background on the implementation of Dawnn. It also provides a brief introduction to the training of neural networks to allow it to be followed by those without a machine learning background.

#### 11.1 Neural Networks

Neural networks are machine learning models [23, 24] capable of approximating any function [25]. They have been shown to be highly effective in a range of both classification and regression problems [26].

A basic *feed-forward neural network* consists of a sequence of *layers*, each of which contains a collection of *nodes*. Commonly (and as is the case with Dawnn), every node in a layer is connected to every node in the layer following it by an *edge*. Two layers each consisting of 1,000 nodes will therefore have 1,000,000 edges connecting them. Each edge is associated with a numerical *weight*, which approximately signifying its importance within the model. The input (in our case, the labels of the 1,000 nearest neighbours of the cell) is passed layer-by-layer through this network, with each node in a layer taking as input the values at all nodes connected to it in the previous layer. The sum of a node’s inputs is then passed through a non-linear function (in our case, the function ReLU(*x*) = max(0*, x*)) and is finally passed along its outgoing edges, with each edge’s value being multiplied by its associated weight. A specific neural network architecture is defined by the number of layers, number of nodes in each layer, and configuration of edges connecting the nodes.

#### 11.2 Training a neural network

It is common to train a neural network using the *backpropagation* algorithm [23]. This approach optimises the edge weights of a model to minimise the difference between its current output and the desired values. A neural network has a number of *hyperparameters* that can affect performance, such as its architecture, learning rate (amount by which weights are updated during training), and whether to employ dropout (where edges are deactivated at random during training to prevent overfitting the training data). For five network architectures, we assessed the performance of two learning rates (0.0001 and 0.00001) and the presence of dropout using five-fold cross validation. Cross validation estimates which neural network architectures will best predict 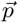 in unseen data by training it five times (*folds*), each time on a different 80% of the training data with the remaining 20% used to assess the learned model. Early stopping [27] was employed in each fold. This reduces the risk of overfitting to the training data by halting training once the performance of the model has not improved for some number of training iterations a separate dataset. We terminated the training process after 25 iterations with no improvement.

#### 11.3 Neural network implementation

The neural network at the heart of Dawnn was implemented in the Python libraries Keras [28] and Tensorflow [29]. The architecture selection process resulted in a choice of 1,000 input notes followed by seven dense layers (i.e. where each node in a layer is connected to every node in the preceding one), with these hidden layers containing 2,000, 4,000, 2,000, 1,000, 250, and 10 nodes. We used the ReLU activation function in each layer except the final one (consisting of a single output node), where we use the sigmoid function to constrain the probabilities to (0, 1). Mean squared error was used as the performance metric and Adam was used as the optimiser [30]. To improve generalisation and training efficiency, we used early stopping with a patience of 25 epochs (i.e. training was halted after 25 epochs in which the mean squared error did not decrease). The code used for training Dawnn’s neural network is available online^7^.

### 12 Proportion-based representation can lose information

To observe that a label-based representation of a dataset loses no information compared with a proportion-based one, consider the vectors [*S_a_ S_b_ S_a_ S_b_*] and [*S_b_ S_a_ S_b_ S_a_*], which represent different *p⃗*s of neighbouring cells in a setting with two labels *S_a_*and *S_b_*.

Note that both vectors of the proportions of the label S*_a_* ([1 0.5 0.667 0.5] and [0 0.5 0.333 0.5], respectively) are identical at distances 2 and 4. Thus the model inputs at these distances would be identical under a proportion-based representation, whereas they differ under a label-based one. Information can thus be lost when using a proportion-based representation.

### 13 Parameter values in benchmarking

The tables below contain the values of the parameters used by each algorithm in the benchmarking procedures. The parameter values used by Dann *et al.* [1] for the mouse dataset were used for the other biological datasets since they are of comparable complexity.

**Supplementary Table 7:** Parameter values of Milo during benchmarking. CNA used the same values of K.

| Dataset | K | d | prop |
| --- | --- | --- | --- |
| Simulated discrete clusters | 15 | 30 | 0.1 |
| Simulated linear trajectory | 20 | 30 | 0.1 |
| Simulated branching trajectory | 20 | 30 | 0.1 |
| Mouse | 50 | 30 | 0.1 |
| Skin | 50 | 30 | 0.1 |
| Organoid | 50 | 30 | 0.1 |
| Heart | 50 | 30 | 0.1 |

**Supplementary Table 8:** Parameter values of DA-seq during benchmarking.

| Dataset | K vector |
| --- | --- |
| Simulated discrete clusters | 15, 65, 115, ..., 465 |
| Simulated linear trajectory | 20, 70, 120, ..., 470 |
| Simulated branching trajectory | 50, 100, 150, ..., 500 |
| Mouse | 50, 100, 150, ..., 500 |
| Skin | 50, 100, 150, ..., 500 |
| Organoid | 50, 100, 150, ..., 500 |
| Heart | 50, 100, 150, ..., 500 |

**Supplementary Table 9:** Parameter values of Dawnn during benchmarking.

| Dataset | $\alpha$ | k |
| --- | --- | --- |
| All | 0.1 | 1,000 |

## Footnotes

1 https://cran.r-project.org/web/packages/dawnn/index.html

2 https://rdr.ucl.ac.uk/projects/Dawnn_manuscript_files/161707

3 https://doi.org/10.5522/04/22633606

4 https://github.com/george-hall-ucl/dawnn_paper_resources

5 https://github.com/george-hall-ucl/dawnn

6 https://github.com/MarioniLab/milo_analysis_2020

7 https://doi.org/10.5522/04/22633606

## Notes

https://github.com/george-hall-ucl/dawnn_paper_resources

### Competing Interest Statement

The authors have declared no competing interest.

### Summary of Updates

Extends the original manuscript to cover local and global definitions of differential abundance.

